# Endothelial SIRPα signaling controls thymic progenitor homing for T cell regeneration and antitumor immunity

**DOI:** 10.1101/2021.04.19.440464

**Authors:** Boyang Ren, Huan Xia, Yijun Liao, Hang Zhou, Zhongnan Wang, Yaoyao Shi, Mingzhao Zhu

**Author notes:** Correspondence author: Mingzhao Zhu.

## Abstract

Thymic homing of hematopoietic progenitor cells (HPCs) is an essential step for the subsequent T cell development. Previously we have identified a subset of specialized thymic portal endothelial cells (TPECs), which is important for thymic HPC homing. However, the underlying molecular mechanism remains still unknown. Here we found that signal regulatory protein alpha (SIRPα) is preferentially expressed on TPECs. Disruption of CD47-SIRPα signaling in mice resulted in reduced number of thymic early T cell progenitors (ETPs) and impaired thymic HPC homing. Mechanistically, SIRPα-deficient ECs and CD47-deficient lymphocytes demonstrated impaired transendothelial migration (TEM). Specifically, SIRPα intracellular ITIM motif-initiated downstream signaling in ECs was found to be required for TEM in a SHP2- and Src-dependent manner. Furthermore, CD47-signaling from migrating cells and SIRPα intracellular signaling were found to be required for VE-cadherin endocytosis in ECs. Functionally, SIRPα signaling is required for T cell regeneration upon sub-lethal total body irradiation (SL-TBI); CD47-SIRPα signaling blockade post SL-TBI diminishes antitumor immunity. Thus, our study reveals a novel role of endothelial SIRPα signaling for thymic HPC homing for T cell regeneration and antitumor immunity.

**Graphic abstract:** 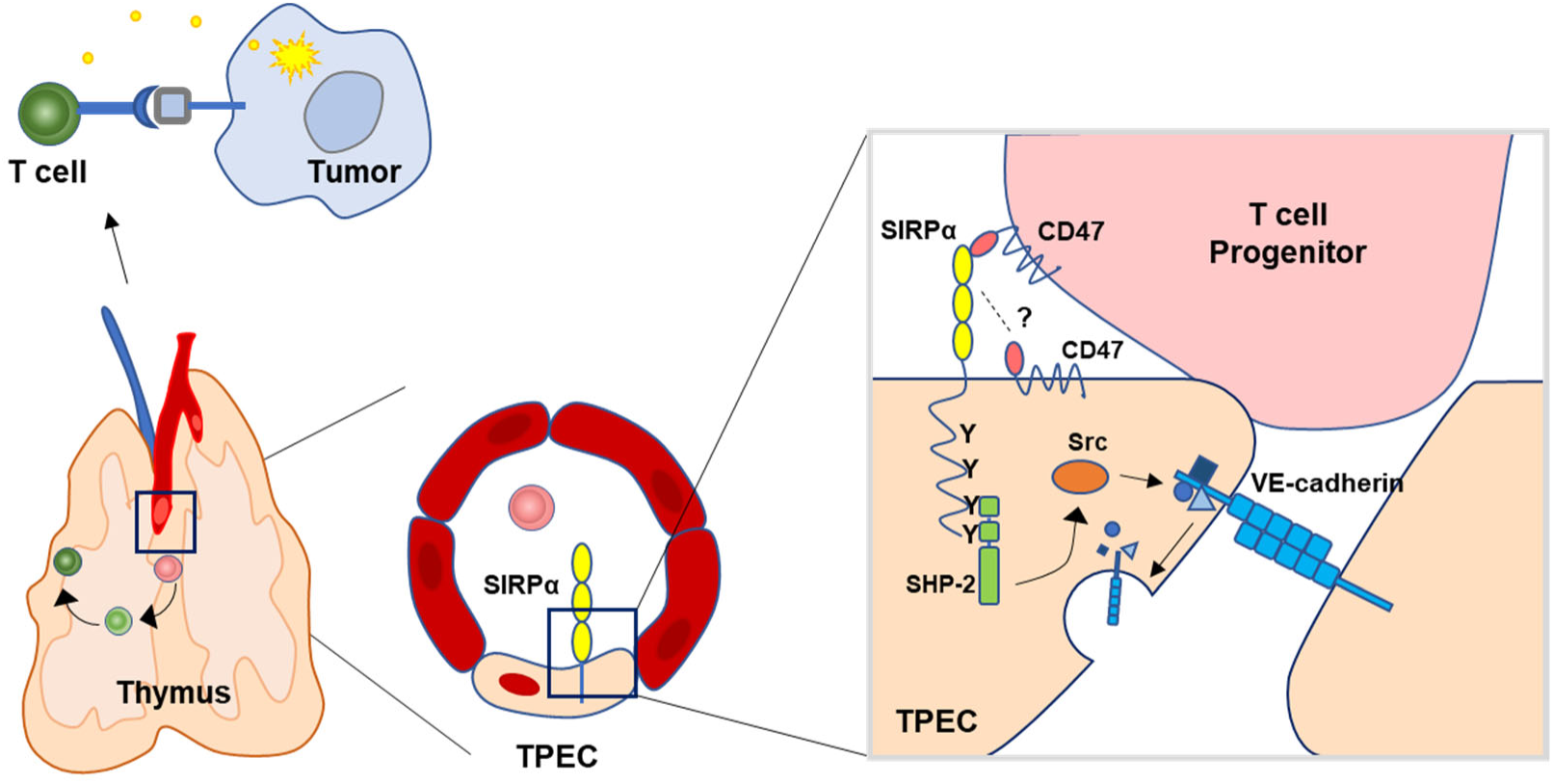

Thymic homing of hematopoietic progenitor cells is fundamental to the T cell-based adaptive immunity, yet the molecular basis of this process is less clear. We discovered that endothelial SIRPα signaling, engaged by migrating cell derived CD47 ligand, regulates thymic homing of hematopoietic progenitor cells for T cell regeneration and antitumor immunity.

- SIRPα is preferentially expressed on thymic portal endothelial cells.
- Endothelial SIRPα regulates thymic homing of hematopoietic progenitor cells.
- CD47-SIRPα downstream signaling induces VE-cadherin endocytosis.
- CD47-SIRPα signaling blockade impairs thymic T cell regeneration and antitumor immunity.

## Introduction

Different from most other hematopoietic cells, T cells develop in the thymus. Thymic homing of bone marrow derived hematopoietic progenitor cells (HPCs) is therefore a critical step. It was reported that HPCs enter the thymus via unique blood vessels that are surrounded by perivascular spaces (PVS)(Lind et al., 2001; Mori et al., 2007) primarily in the corticomedullary junction (CMJ) of the thymus. To understand how the unique blood vascular endothelial cells (ECs) are involved in thymic homing of HPCs is an important question in the field. Previously, we identified a specialized subset of P-selectin^+^ Ly6C^-^ thymic portal endothelial cells (TPECs) located in the CMJ region and associated with PVS structure, providing the cellular basis for thymic homing of HPCs(Shi et al., 2016). Short-term HPC thymic homing assay confirmed that TPECs is highly associated with settling HPCs and lack of TPECs in lymphotoxin beta receptor deficient mice resulted in dramatically impaired HPC thymic homing and reduced number of thymic early T cell progenitors (ETPs). Transcriptome analysis revealed that TPECs are enriched with transcripts related to cell adhesion and trafficking, supporting its critical role for thymic homing. However, the molecular basis of TPEC function and its underlying mechanism have not been experimentally determined.

Several molecules have been found to play important roles for thymic homing of HPCs. P-selectin and adhesion molecule VCAM-1 and ICAM-1, highly expressed on ECs, as also confirmed in our previous study, have been suggested to mediate adhesion of HPCs on ECs(Lind et al., 2001; Mori et al., 2007; Rossi et al., 2005; Scimone et al., 2006). In addition, chemokines such as CCL19 and CCL25 expressed by thymic ECs as well as thymic epithelial cells (TECs) are also involved in thymic homing of HPCs, probably via integrin activation(Krueger et al., 2010; Misslitz et al., 2004; Parmo-Cabañas et al., 2007; Zhang et al., 2014; Zlotoff et al., 2010). All the above-mentioned mechanisms regulate the multistep HPC homing at early invertible process(Zlotoff and Bhandoola, 2011). Transendothelial migration (TEM) is the decisive step for migration of progenitors from blood into the thymus(Zlotoff and Bhandoola, 2011). To accomplish the homing process, the barrier of endothelial junction must be diminished during migration. How this step is regulated in TPECs remains still unknown.

Signal regulatory protein alpha (SIRPα) is a transmembrane protein that contains three Ig-like domains, a single transmembrane region, and a cytoplasmic region. Ligation of SIRPα by its ligand CD47 transmits intracellular signal through its ITIM motifs. SIRPα is mainly expressed by myeloid cells such as monocytes, granulocytes, most tissue macrophages and subsets of dendritic cells(Barclay and Van den Berg, 2014). On the other hand, CD47 is ubiquitously expressed but show fluctuating expression levels on different cell states or cell types. Elevated CD47 expression is detected on bone marrow cells and some of lymphoid subsets(Jaiswal et al., 2009; Van et al., 2012).

SIRPα plays various roles in immune system. SIRPα on conventional dendritic cells (cDCs) maintains their survival and proper function via intracellular SHP2 signaling(Iwamura et al., 2011; Saito et al., 2010). On macrophages, upon CD47 ligation, SIRPα signaling activates intracellular SHP1 to inhibit Fcγ receptor-mediated phagocytosis towards target cells(Blazar et al., 2001; Ishikawa-Sekigami et al., 2006; Tsai and Discher, 2008). Elevated expression of CD47 on platelets, lymphocyte subsets and hematopoietic cell subsets shows ‘self’ identity and protects them from being cleared by macrophages(Blazar et al., 2001; Jaiswal et al., 2009; Olsson et al., 2005; Yamao et al., 2002). Tumor cells express high level of CD47 to escape immune surveillance initiated by macrophages(Chao et al., 2010; Majeti et al., 2009; Willingham et al., 2012), which leads to the development of CD47-SIRPα blockade as a promising approach for cancer therapy.

CD47-SIRPα has also been reported to play important roles in cell adhesion and migration. SIRPα expressed on cDCs, neutrophils, melanoma cells and CHO cells has been reported to promote cell motility(Fukunaga et al., 2004; Liu et al., 2002; Motegi et al., 2003) through SHP2-dependent activation of Rho GTPase and cytoskeleton reorganization(Wollenberg et al., 1996),(Inagaki et al., 2000). On the other hand, SIRPα expressed on neutrophils and monocytes has been shown to be the ligand for endothelial or epithelial CD47 and promotes junction opening through activating Rho family GTPase therefore permitting transmigration of the SIRPα-expressing cells(de Vries et al., 2002; Liu et al., 2002; Stefanidakis et al., 2008). However, the expression and function of SIRPα on ECs remain unknown.

In the present study, we uncovered the novel role of TPEC signature molecule SIRPα in thymic homing of progenitor cells, revealed that migrating cell-derived CD47 and EC-SIRPα intracellular signal induce junctional VE-cadherin endocytosis and promote TEM. In addition, we showed that CD47-SIRPα signaling blockade upon thymic injury resulted in impaired T cell regeneration and antitumor immunity.

## Results

### Thymic portal endothelial cells preferentially express SIRPα

To explore the molecular mechanisms for thymic homing of the progenitors through TPECs, we started with signature genes of TPECs based on previously published RNA-seq data and focused on the genes related to cell adhesion and migration by intersecting with related gene sets(Shi et al., 2016)(**SupplementaryFile1)**. Among those signature genes of TPECs, SIRPα (**Figure 1A,B**) is of particular interest since its involvement in cell migration on leukocytes but undiscovered role on non-immune cells.

**Figure 1.**
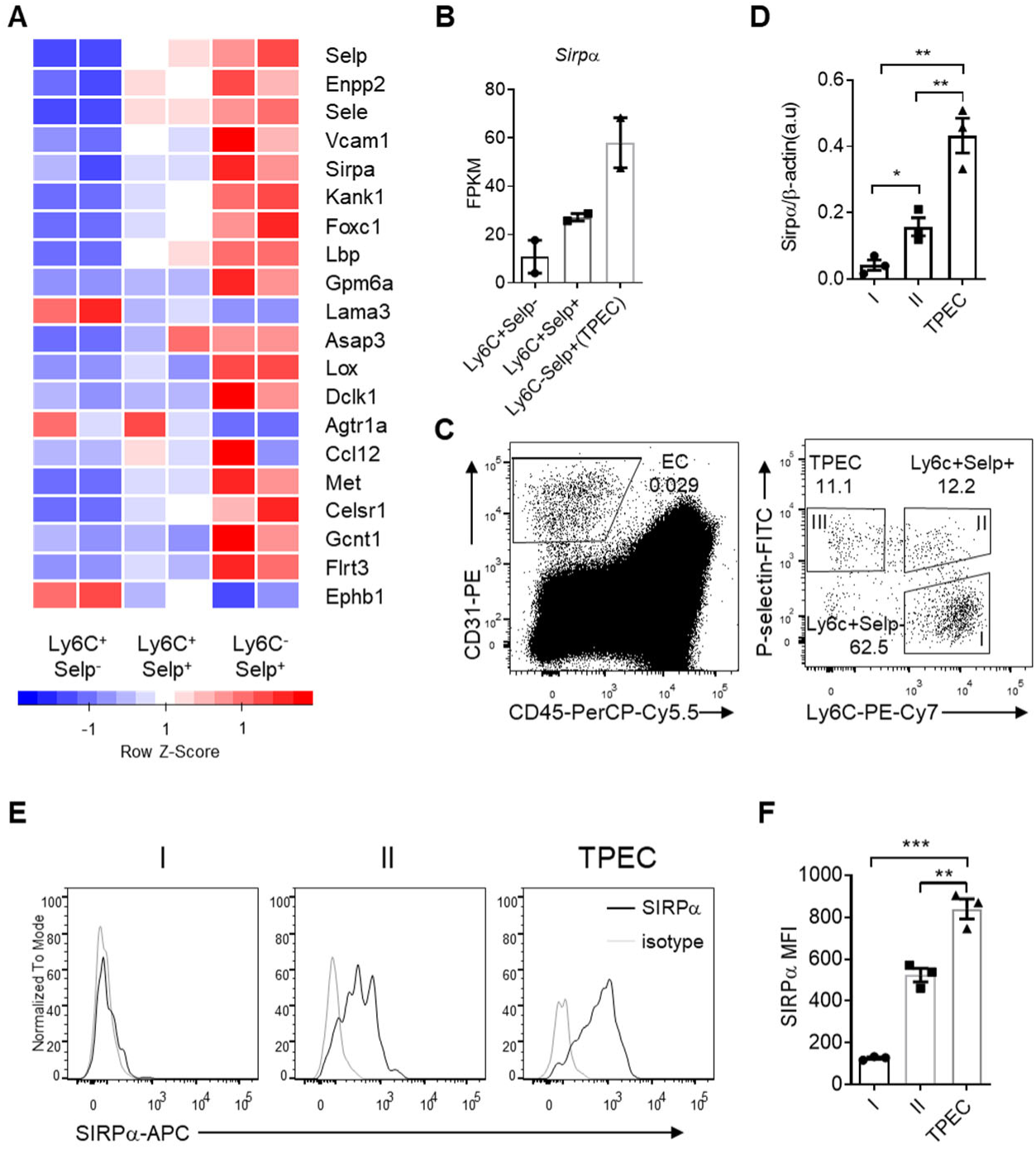
SIRPα is preferentially expressed on TPECs. (**A**) Expression profile of top 20 signature genes of Ly6C^-^Selp^+^ ECs (TPEC), which have absolute FC>2 and p<0.01 in TPECs versus either Ly6C^+^Selp^-^ or Ly6C^+^Selp^+^ thymic EC subsets, and are in GO term GO_0016477 (cell migration). Relative expression of each gene among EC subsets are presented as mean-centered z-score distribution. (**B**) Expression level of *Sirpα* among the three thymic EC subsets. FPKM: fragments per kilobase per million mapped reads. (**C**) Flow cytometric analysis of thymic ECs (CD31^+^CD45^-^) and subset I (Ly6C^+^Selp^-^), subset II (Ly6C^+^Selp^+^) and subset III (TPEC, Ly6C^-^Selp^+^). (**D**) Real-time PCR analysis of *Sirpα* mRNA expression in thymic EC subsets. (**E-F**) Flow cytometry analysis of SIRPα expression on the three thymic EC subsets (**E**) and quantification of measuring mean fluorescence intensity of SIRPα (**F**), data are representative of three independent experiments with three mice in each experiment. Error bars represent s.e.m. Asterisks mark statistically significant difference, \**P*<0.05, \*\**P*<0.01 and \*\*\**P*<0.001 determined by two-tailed unpaired Student’s *t*-test. Source data and detailed method for generating heatmap in A. are available in SupplementaryFile1.

To further assess the expression of SIRPα on thymic endothelial subsets, CD31^+^ thymic ECs were further separated by Ly6C and P-Selectin (**Figure 1C**) as previously reported(Shi et al., 2016), and analyzed by quantitative RT-PCR and flow cytometry. SIRPα was barely detectable on Ly6C^+^P-Selectin^-^ ECs, the dominant population of thymic ECs. On Ly6C^+^P-Selectin^+^ ECs, the suggested precursor of TPECs, there was a substantial level of SIRPα expression. Among all three EC subsets, Ly6C^-^P-Selectin^+^ TPECs, express the highest level of SIRPα at both mRNA level (**Figure 1D**) and protein level (**Figure 1E,F**). Thus, SIRPα appears to be closely and positively related to thymic EC portal function.

### Endothelial SIRPα is essential for ETP population maintenance and thymic progenitor homing

ETPs are the very first subset of bone marrow-derived progenitor cells that settle the thymus and are committed to T cell lineage. Studies have suggested ETP population size a direct indicator of thymic HPC homing(Krueger et al., 2010; Shi et al., 2016; Zlotoff et al., 2010). Therefore, to test the requirement of SIRPα for thymic progenitor homing, we first analyzed ETP population in *Sirpα*^-/-^ mice. *Sirpα*^-/-^ mice appeared normal regarding to the total cellularity in thymus (**Figure 2—figure supplement 1A**) and peripheral lymphoid organs (**Figure 2—figure supplement 1B,C**). The frequencies and numbers of major subsets of thymocytes remain unaltered except for DN1 subset which mainly consists of ETPs (**Figure 2—figure supplement 1D-G**). A substantial reduction in ETP population was confirmed in *Sirpα*^-/-^ mice (**Figure 2A,B**). Bone marrow multipotent progenitors including lineage^-^Sca-1^+^c-Kit^high^ (LSK) and common lymphoid progenitors (CLPs) remained unchanged in these mice (**Figure 2C,D**), suggesting the reduced population size of thymic ETP is unlikely due to impaired generation/homeostasis of bone marrow progenitors.

**Figure 2.**
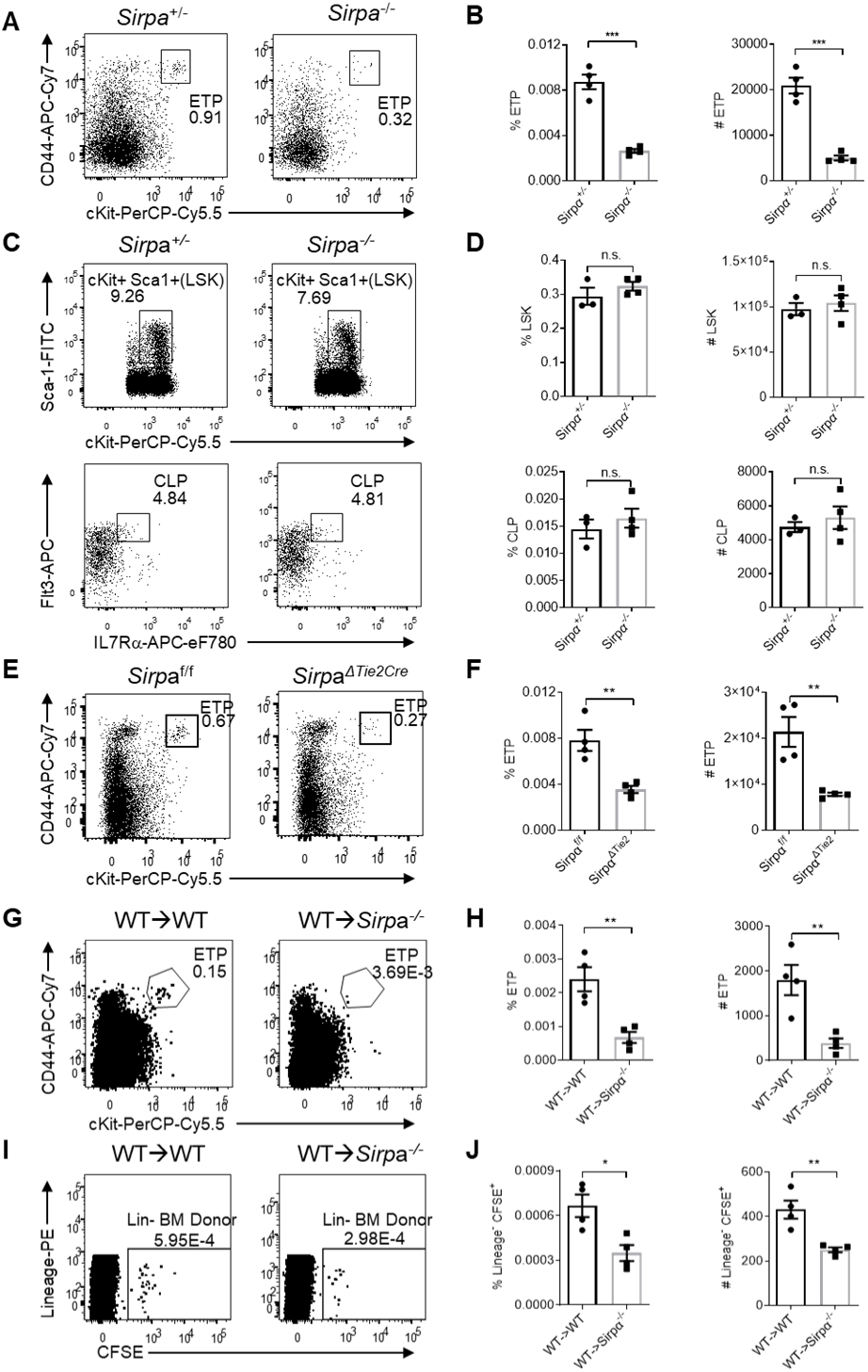
Endothelial SIRPα is essential for ETP population maintenance and thymic progenitor homing. (**A**) Representative flow cytometric analysis of ETPs (Lineage^-^CD25^-^CD44^+^cKit^+^) in the thymus of *Sirpα^-/-^* and control mice. (**B**) Proportion of ETP population of total thymocytes and corresponding cell number in a thymus. (**C**) Analysis of LSKs (Lineage^-^Sca1^+^cKit^+^) and CLPs (Lineage^-^Sca1^lo^cKit^lo^Flt3^+^IL7Rα^+^) in the bone marrow. (**D**) Proportion and total cell number of LSK and CLP in a pair of femurs and tibias. (**E**) Flow cytometric analysis of ETPs in *Sirpα*^Δ^*^Tie2Cre^* conditional knock out mice. (**F**) Statistics of ETPs in the thymus. (**G**) Representative flow cytometric analysis of ETPs (Lineage^-^CD25^-^CD44^+^cKit^+^) in WT bone marrow reconstituted WT or Sirpα^-/-^ mice. (**H**) Statistics of ETPs in the thymus. (**I,J**) Short-term homing assay in WT bone marrow reconstituted WT or Sirpα^-/-^mice. (**I**) Analysis of lineage-negative donor cells (Lineage^-^ CFSE^+^) cells among total thymocytes. (**J**) Statistics of lineage-negative donor cells in the thymus. Data are representative of three independent experiments with three mice in each experiment. Error bars represent s.e.m. Asterisks mark statistically significant difference, \**P*<0.05, \*\*\**P*<0.001, \*\*\*\**P*<0.0001, n.s. not significant, determined by two-tailed unpaired Student’s *t*-test.

Previous studies suggested that thymic entry of the progenitor cells and ETP population maintenance is guided by multiple cues derived from thymic endothelial cells as well as thymic epithelial cells. To specifically test the role of endothelial SIRPα, we generated *Sirpα*^ΔTie2-Cre^ (*Sirpα*^flox/flox^ × Tie2-cre) mice. Significant decrease of thymic ETP population was found in *Sirpα*^ΔTie2-Cre^ mice in a degree similar to that in *Sirpα*^-/-^ mice (**Figure 2E,F**), suggesting the role of SIRPα is probably mainly derived from ECs. In supporting this, much lower expression level of SIRPα was found on either medullary or cortical thymic epithelial cells (**Figure 2—figure supplement 2A-C**).

Previous studies have reported SIRPα as a phagocytic checkpoint on tissue macrophages and CD47-null cells are rapidly cleared in congenic wild type (WT) mice(Bian et al., 2016). Possibility exists that phagocytic activity may elevate towards circulating progenitor cells in *Sirpα*^-/-^ or *Sirpα*^ΔTie2-Cre^ mice before they reach the thymus, since SIRPα is deficient in hematopoietic cells of both mouse lines. To distinguish the role of hematopoietic cell-derived versus non-hematopoietic radioresistant stromal cell-derived SIRPα in the regulation of thymic ETPs, bone marrow chimeric mice were generated (**Figure 2—figure supplement 2D**). Mice lacking SIRPα on radioresistant stromal cells demonstrated significantly reduced ETP population (**Figure 2G,H**), whereas in mice lacking SIRPα on hematopoietic cells, ETP population remained unchanged (**Figure 2—figure supplement 2E,F**). Thus, together with previous data, reduced thymic ETP population in *Sirpα*^-/-^ or *Sirpα*^ΔTie2-Cre^ mice is unlikely due to increased clearance of HPCs in the absence of hematopoietic SIRPα.

To directly test the role of SIRPα on HPC thymic homing, a short-term thymic homing assay was adopted(Shi et al., 2016), in which massive amount of progenitor-containing bone marrow cells were intravenously transferred and thymic settling progenitor cells were determined two days later. To separate the hematopoietic versus non-hematopoietic role of SIRPα, bone marrow chimeric mice was used for short-term homing assay (**Figure 2—figure supplement 2D**). SIRPα deficiency on hematopoietic cells did not result in impaired thymic settling of bone marrow progenitors (**Figure 2—figure supplement 2G**), nor did the number of progenitors retained in the periphery as represented by the number of progenitor cells in the spleen (**Figure 2—figure supplement 2H**). On the contrary, loss of SIRPα on radioresistant cells significantly impeded thymic entry of progenitor cells (**Figure 2I,J**). Nevertheless, unaltered number of donor bone marrow cells retained in the periphery (**Figure 2—figure supplement 2I,J**), further confirming the role of endothelial SIRPα in controlling thymic entry of the progenitors. Together, these data suggest that myeloid-SIRPα mediated phagocytic activity does not affect thymic progenitor homing in the current system; endothelial SIRPα plays an important role on thymic homing of progenitor cells and ETP population maintenance.

### EC-SIRPα controls lymphocyte TEM

Since TPECs is the portal of thymic progenitor entry, we asked whether SIRPα signaling may regulate TPEC development. *Sirpα*^-/-^ mice did not show altered thymic endothelial development in regard to the percentage and number of total ECs (**Figure 3—figure supplement 1A-C**) and specifically TPECs (**Figure 3A-C**). Adhesion of progenitor cells to the wall of blood vessel is an early step for successful transmigration towards thymic parenchyma. Since SIRPα has been reported for its role in cellular adhesion(Seiffert et al., 1999), we next tested the key adhesion molecules involved in leukocyte-EC adhesion and found them largely unchanged on TPECs of *Sirpα*^-/-^ mice (**Figure 3D,E**). Concerning other potential effect of SIRPα on leukocyte-EC adhesion, we directly tested this *in vitro*. *Sirpα*^-/-^ MS1 endothelial cell line was constructed by CRISPR-Cas9 and the deletion of SIRPα was confirmed by flow cytometry (**Figure 3—figure supplement 1D**). Compared to WT ECs, *Sirpα*^-/-^ EC monolayer mediates comparable adhesion of lymphocytes (**Figure 3F**).

**Figure 3.**
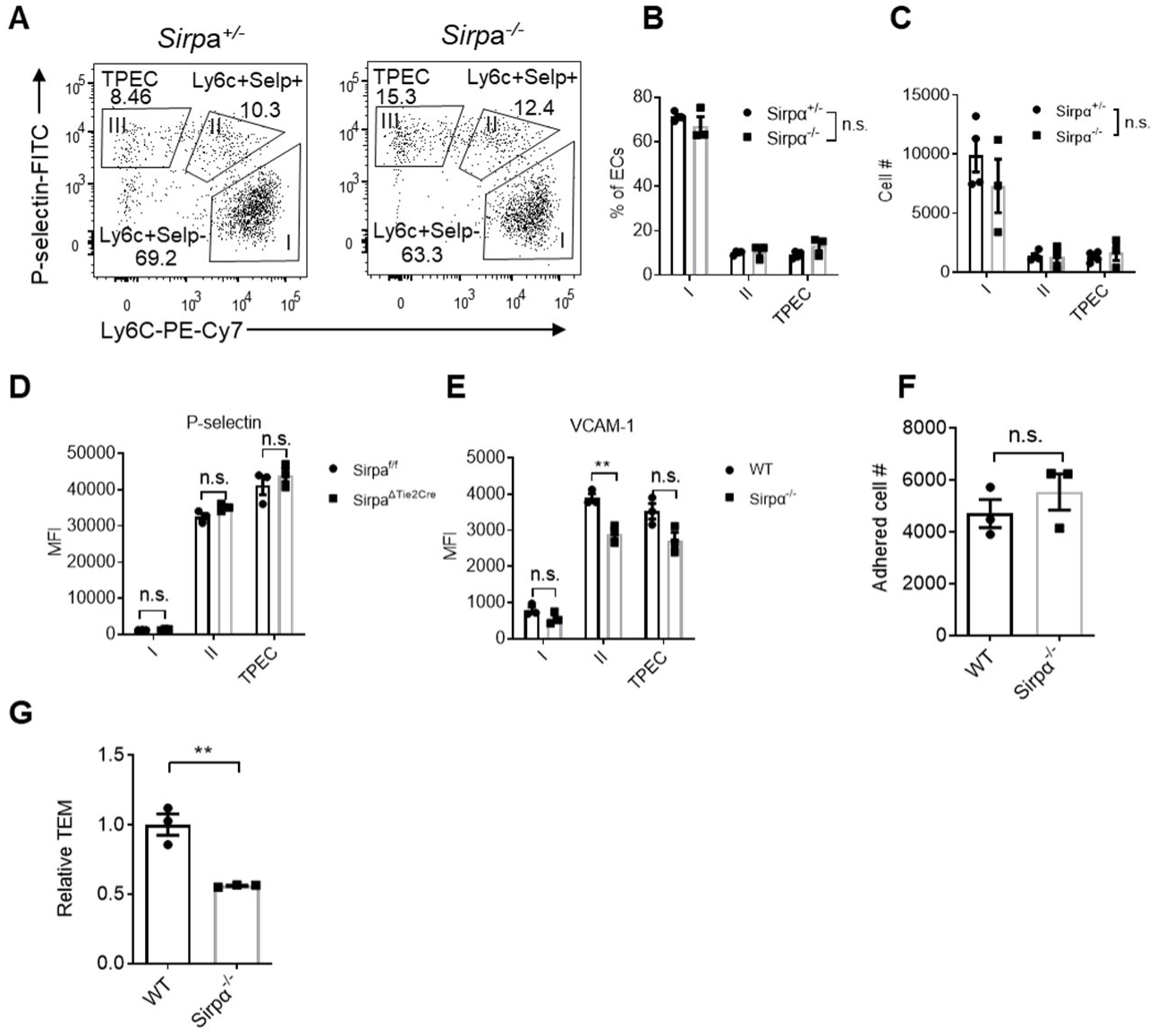
EC-SIRPα controls lymphocyte TEM. (**A**) Representative flow cytometric analysis of thymic EC composition in *Sirpα^-/-^* and control mice. (**B,C**) Proportion of thymic EC subsets (**B**) and corresponding cell numbers (**C**) in the thymus. (**D,E**) Expression level of adhesion molecules on thymic EC subsets. (**D**) P-selectin in *Sirpα*^Δ^*^Tie2Cre^* or control mice. (**E**) VCAM-1 in *Sirpα^-/-^* or control mice. (**F**) Adhered lymphocyte cell number in a well in the cell adhesion assay on *Sirpα^-/-^* or WT MS1 endothelial cells. (**G**) Lymphocyte transmigration in the presence of CCL-19 (10ng/ml) in the lower chamber of transwell, quantified by flow cytometry. Transmigrated cell number in the lower chamber in WT group was normalized to 1. Data information: Error bars represent s.e.m. Asterisks mark statistically significant difference, \*\**P*<0.01, n.s. not significant, determined by two-tailed unpaired Student’s *t*-test. Raw data of transmigration assay in G are available in SupplementaryFile1.

Next, we examined the role of SIRPα in TEM by transwell assay. Monolayers of both WT and *Sirpα*^-/-^ ECs were equally formed at the time of transmigration assay (**Figure 3—figure supplement 1E,F**). In the presence of CCL19 in the lower chambers, T lymphocytes were measured for their transmigration across endothelial barrier. Remarkably, SIRPα deficiency in ECs resulted in about 45% reduction of transmigrated cell numbers (**Figure 3G**), suggesting a direct role of SIRPα in controlling the behavior of ECs in the process of lymphocyte TEM.

### Migrating cell-derived CD47 guides their TEM

CD47, the cellular ligand of SIRPα, is ubiquitously expressed on almost all types of cells. Interestingly, we found among developmental subsets of T cell lineage, ETPs exhibited the highest level of CD47 expression (**Figure 4A**), suggesting preferential interaction may exist between ETPs and TPECs controlling thymic progenitor homing and ETP population. Indeed, *Cd47^-^*^/-^ mice showed significant reduced ETP population (**Figure 4B-D**) and unaffected ancestral progenitor subsets (**Figure 4—figure supplement 1A-C**), similar to that of *Sirpα*^-/-^ mice. To directly test the role of CD47 on migrating cells in TEM, the transwell assay was applied as described above. Significantly elevated CD47 expression was found on immortalized cells, herein MS1 (**Figure 4—figure supplement 1D**), which was not found in the physiological situation, wherein EC-CD47 expression level is much lower than that on migrating cells (ETP) (**Figure 4—figure supplement 1E,F**). To exclude the potential influence of this artificial CD47 expression on ECs and to determine the sole role of migrating cell-derived CD47, *Cd47^-/-^* MS1 cell line was generated (**Figure 4—figure supplement 1G**). Either genetic deficiency of CD47 or blockade of CD47 signal on lymphocytes via SIRPα-hIg (CV-1) resulted in significantly fewer TEM through *Cd47^-/-^* MS1 cells (**Figure 4E,F**). Thus, CD47 expression on lymphocyte correlated with enhanced transmigration. These data suggest that elevated CD47 on migrating progenitors could engage SIRPα on specialized TPECs to promote their thymic entry.

**Figure 4.**
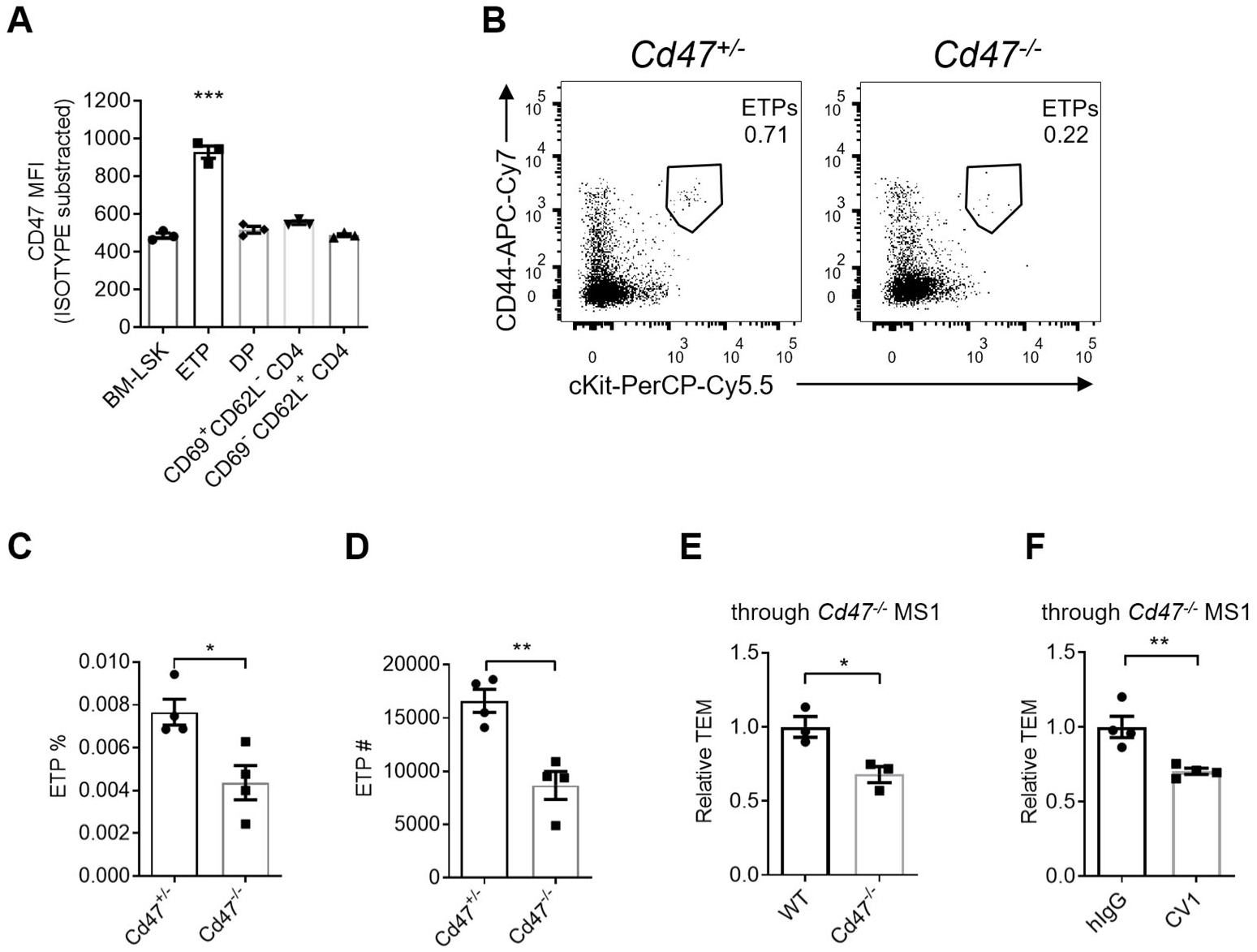
Migrating cell-derived CD47 guides their TEM. (**A**) CD47 expression measured by flow cytometry in various subsets of T cell lineage. BM-LSK: Lineage^-^Sca1^+^cKit^+^ lymphoid progenitor cells in bone marrows; ETP: early T-cell progenitors in the thymus; DP: double positive (CD4^+^CD8^+^) thymocytes; CD69^+^CD62L^-^CD4^+^ and CD69^-^CD62L^+^CD4^+^: immature and mature CD4 single positive thymocytes, respectively. (**B**) Flow cytometric analysis of ETPs. (**C,D**) Proportion (**C**) and corresponding cell numbers (**D**) of ETP population in a thymus in *Cd47^-/-^* or control mice. (**E**) Transmigrated *Cd47^-/-^* or WT lymphocytes through *Cd47^-/-^* MS1 endothelial monolayer. (**F**) Transmigrated CV1-incubated or control lymphocytes through *Cd47^-/-^* MS1 endothelial monolayer. Lymphocytes were incubated with CV1 (10μg/ml) or control hIg (10μg/ml) for 30 minutes at 4℃ before applied to the transwell. Error bars represent s.e.m. Asterisks mark statistically significant difference, \**P*<0.05, \*\**P*<0.01, \*\*\**P*<0.001, determined by two-tailed unpaired Student’s *t*-test. Raw data of transmigration assay in E and F are available in SupplementaryFile1.

### SIRPα intracellular signal controls TEM via SHP2 and Src

CD47-SIRPα interaction could transduce signaling bidirectionally. We next examined whether SIRPα signals through ECs for TEM regulation. To do this, an endothelial cell line lacking intracellular domain of SIRPα was constructed(Inagaki et al., 2000) (SIRPα-ΔICD MS1) (**Figure 5—figure supplement 1A**). SIRPα-ΔICD MS1 cells did not abolish surface display of the molecule and considerable expression of extracellular region of SIRPα is detected (**Figure 5—figure supplement 1B**), which would retain its ability to engage CD47 for potential signaling downstream of CD47 in interacting cells. However, lymphocyte transmigration across SIRPα-ΔICD MS1 cells reduced to the level similar to that on *Sirpα^-/-^* MS1 cells (−33% of WT, comparing −42% in *Sirpα^-/-^*) (**Figure 5A**), suggesting SIRPα regulates TEM mainly through ECs. Furthermore, lentiviral transduction with WT *Sirpα* coding sequence (*Sirpα*-WT) in *Sirpα^-/-^* MS1 cells largely recovered TEM (78% of WT level, comparing 32% of WT in *Sirpα^-/-^* MS1) (**Figure 5B**), whereas transduction with loss-of-function mutant *Sirpα*-Y4F, in which all four functional tyrosine residues in the cytoplasmic ITIM motifs were substituted with inactive phenylalanine, failed in TEM rescue (23% of WT) (**Figure 5B**), even with restored surface expression (**Figure 5—figure supplement 1C**). Thus, SIRPα intracellular ITIMs-mediated downstream signaling in ECs is required for TEM.

**Figure 5.**
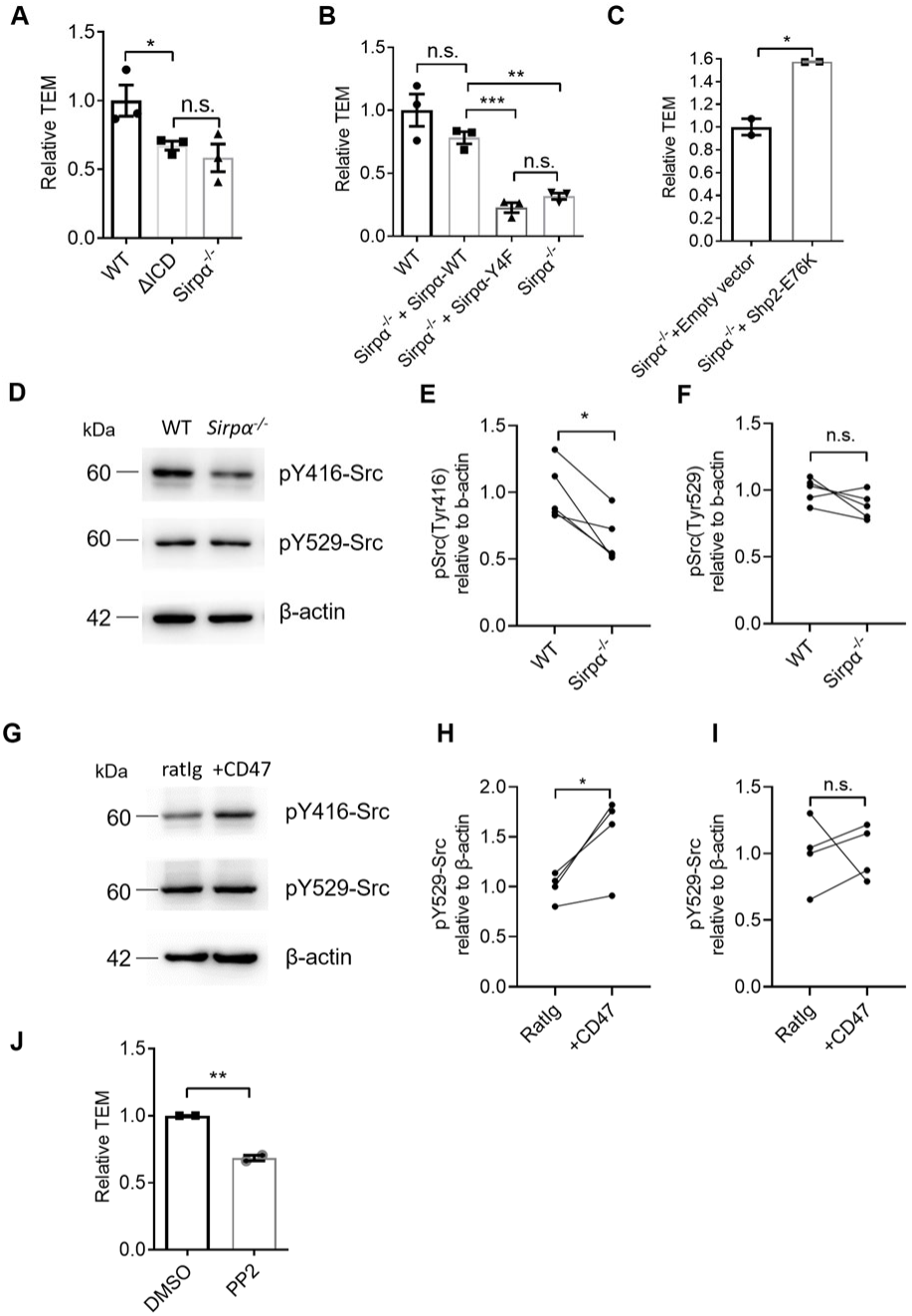
SIRPα intracellular signal controls TEM via SHP2 and Src. (**A**) Relative TEM efficiency of lymphocytes through WT, intracellular-truncated SIRPα-ΔICD, and *Sirpα^-/-^* (KO) MS1 endothelial monolayer. (**B**) Relative TEM efficiency of lymphocytes through *Sirpα* overexpressing (*Sirpα^-/-^*+Sirpα-WT) or Sirp*α*-tyrosine-to-phenylalanine mutant overexpressing (*Sirpα^-/-^*+Sirpα-4F) MS1 endothelial monolayer. **(C)** Relative TEM efficiency of lymphocytes through *Sirpα^-/-^* MS1 endothelial monolayer overexpressing constitutively active form of SHP2 (+*Shp2*-CA) or control empty vector (+ Empty vector). (**D**) Detection of Src activity in *Sirpα^-/-^* and WT MS1 endothelial cell lines, as measured by western blot with anti-pY416-Src (active form Src antibody), anti-pY529-Src (inactive form Src antibody) and anti-β-actin control antibody. The plot is representative of five independent experiments. (**E,F**) Relative quantification of the active Src (pY416-Src) (**E**) and inactive Src (pY529-Src) (**F**) to β-actin in five tests, the relative expression in each sample is normalized to WT average. (**G**) Detection of Src activity in *Cd47*^-/-^ MS1 cells upon mCD47-Ig or control Ig stimulation, as measured by western blot. The plot is representative of four independent experiments. (**H,I**) Relative quantification of the active Src (pY416-Src) (**H**) and inactive Src (pY529-Src) (**I**) to β-actin in four tests, and the relative expression in each sample is normalized to WT average. (**J**) Lymphocyte transmigration, in the absence or presence of Src inhibitor PP2 (5μM, 12 hours)) through WT MS1 endothelial monolayer. Error bars represent s.e.m. Asterisks mark statistically significant difference, \**P*<0.05, \*\**P*<0.01, \*\*\**P*<0.001, n.s. not significant. Raw data of transmigration assay in A, B, C, J and raw image and data of western blotting in D-I are available in SupplementaryFile1.

Phosphorylation of ITIMs of SIRPα cytoplasmic tail may recruit and activate ubiquitously expressed tyrosine phosphatase SHP2 in ECs(Takada et al., 1998). To investigate whether SHP2 is sufficient to enable TEM in the absence of SIRPα signaling, *Sirpα*^-/-^ MS1 cells were transduced with a lentiviral vector to express constitutively active SHP2 (E76K(Bentires-Alj et al., 2004), hereafter called SHP2-CA). SHP2-CA MS1 cells permitted significantly more (+57% of empty vector control) TEM of lymphocytes (**Figure 5C**), demonstrating that SHP2 activation can rescue the defective effect of SIRPα deficiency.

Several studies have illustrated that SHP2 activity could cooperate activation of Src kinase in ECs(Liu et al., 2012; Zhang et al., 2004), we next investigated whether Src participates TEM regulation in the context of SIRPα signaling. In *Sirpα^-/-^* MS1 cells, active Src (pTyr-416) was significantly deficient compared to WT **(Figure 5D,E)** without affecting the pool of autoinhibitory inactive form of Src **(Figure 5D,F)**. Furthermore, plate-bound CD47 significantly promoted Src activation (pTyr-416,) but not pYyr-529 in *Cd47^-/-^* MS1 cells (**Figure 5G-I**). These data suggest the ability of SIRPα downstream signaling in activating Src kinase.

To directly assess the role of Src in leukocyte transmigration, a specific inhibitor of Src family kinases, PP2, was applied to the WT MS1 cells during transwell assay. Significantly inhibited lymphocyte transmigration was observed (**Figure 5J**). Thus, these data suggest that SIRPα downstream signaling activates SHP2 and Src kinase for the control of TEM.

### CD47-SIRPα signaling promotes VE-cadherin endocytosis

VE-cadherin is the dominant gate-keeping molecule controlling leukocyte transmigrating through endothelial barrier(Corada et al., 1999; Vestweber, 2007; Vestweber et al., 2009; Wessel et al., 2014). VE-cadherin controls junction opening by regulated destabilization and endocytosis from junctional area of cell surface(Allingham et al., 2007; Benn et al., 2016; Wessel et al., 2014). In addition, phosphatase SHP2 and kinase Src have been both reported to destabilize adherens junction during transmigration by promoting endocytosis of VE-cadherin(Allingham et al., 2007; Wessel et al., 2014). To test the role of endothelial SIRPα signal on VE-cadherin endocytosis, we adopted an *in vitro* VE-cadherin endocytosis assay as previously described(Wessel et al., 2014). The surface expression of VE-cadherin is not altered in *Sirpα*^-/-^ ECs (**Figure 6—figure supplement 1A,B**). Upon lymphocyte engagement, *Sirpα^-/-^* MS1 showed significantly lower T-cell induced VE-cadherin endocytosis (**Figure 6A,B**). SIRPα-ΔICD MS1 cells showed similar unresponsiveness of VE-cadherin endocytosis to lymphocyte engagement (**Figure 6C.D**), demonstrating requirement of SIRPα intracellular signaling in facilitating VE-cadherin endocytosis. Furthermore, inhibition of SIRPα downstream molecule Src by PP2 significantly inhibited endocytosis of VE-cadherin in WT MS1 cells but not in *Sirpα^-/-^* MS1 cells (**Figure 6E**), suggesting an important role of Src activity downstream of SIRPα intracellular signaling in controlling VE-cadherin endocytosis.

**Figure 6.**
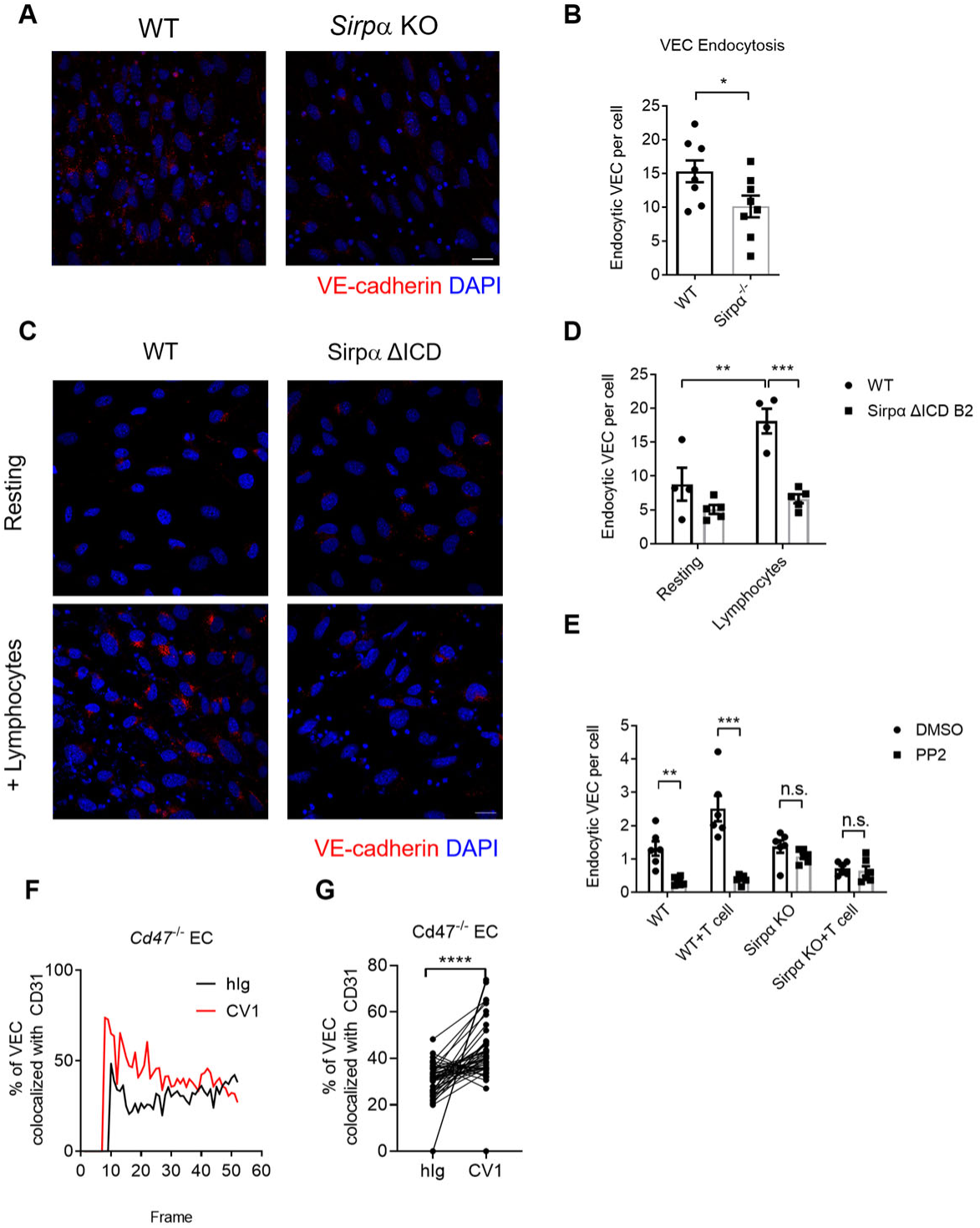
CD47-SIRPα signaling promotes VE-cadherin endocytosis. (**A**) Representative imaging of endocytosed VE-cadherin in the presence of lymphocytes, scale bars represent 20μm. (**B**) Statistical analysis of VE-cadherin endocytosis in MS1 endothelial cells. VE-cadherin fluorescence signal was quantified by ImageJ with same threshold for each slide. MS1 cell number was quantified by DAPI, the final results were presents as arbitrary intracellular VE-cadherin signal per MS1 cell, average signal of all cells was calculated for each filed. Data are representative of three independent experiments with at least five fields (dots on the graph) in each group (captured filed). (**C,D**) VE-cadherin endocytosis in intracellular-truncated SIRPα-ΔICD or WT MS1 cells, with (+Lymphocytes) or without lymphocyte incubation (Resting). Representative confocal imaging of endocytosed VE-cadherin is shown (**C**) with statistical analysis of VE-cadherin endocytosis (**D**). (**E**) VE-cadherin endocytosis in the presence of lymphocytes (+ T cell) and inhibitor of Src activation (+PP2, 5μM, 2 hours). (**F,G**) Real-time analysis of adherens junctional VE-cadherin in the presence of migrating lymphocytes pretreated with CV1 or control hIg. Dynamic measurement (**F**) and statistical analysis of VE-cadherin colocalization with CD31 (**G**) after CD4^+^ T cell injection to the flow. Error bars represent s.e.m. Asterisks mark statistically significant difference, \**P*<0.05, \*\**P*<0.01, \*\*\**P*<0.001, n.s. not significant.Raw data and analysis method of endocytosis assay in A-E and real-time image statistics in F and G are available in SupplementaryFile1.

VE-cadherin endocytosis is a quite dynamic process induced upon cellular contact. In an effort to analyze the effect of CD47-SIRPα signaling on this process, we set up an *in vitro* live imaging procedure to analyze VE-cadherin endocytosis at sites of migrating cell contact(Kroon et al., 2014). CD47-blocked (CV1) or control (hIg treatment) T lymphocytes with different fluorescent labels were mixed equally and then infused into flow chamber mimicking physiological blood flow. The junctional retainment of VE-cadherin was measured by its colocalization with adhesion molecule CD31 (**Figure 6—figure supplement 1C**). VE-cadherin colocalization with CD31 under area of migrating cells was calculated and grouped by their different fluorescent labels. The higher colocalization indicates less VE-cadherin endocytosis. The test was performed on *Cd47*^-/-^ MS1 cells to exclude potential influence of CD47 on ECs. Significantly higher degree of VE-cadherin/CD31 colocalization was found under migrating cells with CD47-SIRPα blockade pretreatment compared to that under control treated migrating cells (**Figure 6F,G**), suggesting the requirement of CD47 signal derived from migrating cells for induction of VE-cadherin endocytosis. Reciprocal fluorescence labeling of CV1- and hIg-treated migrating cells showed similar results (**Figure 6—figure supplement 1D,E**). Together, these data suggest that upon migrating cell contacts, CD47-SIRPα signal regulates VE-cadherin endocytosis via activation of SHP2 and Src kinase, at sites of contact, to control TEM.

### SIRPα is required for thymic regeneration and T cell immune regeneration

Thymic injury occurred in cancer chemoradiotherapy (CRT). Efficient thymic regeneration is in part constrained by the number of progenitors entering thymus(Zlotoff et al., 2011). Given the role of CD47-SIRPα signaling in thymic progenitor homing, we tested whether CD47-SIRPα signal blockade might accidentally impair thymic regeneration. Mice received sub-lethal total-body irradiation (SL-TBI) were intravenously injected with CV1 or control hIg during the period of immune regeneration (**Figure 7A**). SL-TBI caused a dramatic drop of lymphocyte cell numbers in the peripheral blood on day 9 post irradiation(Shi et al., 2016). Thymic progenitors travel the thymus in around 2 weeks, differentiating along their journey, and become mature to be ready to replenish the peripheral T cell pool(Hale and Fink, 2009). T cell regeneration was examined at day 21 post irradiation. CV1-blockade group showed a significant reduction in thymic total cellularity (**Figure 7B**). Flow cytometry analysis showed significantly decreased number of ETP and DP cells in the thymus of CV1-treated mice (**Figure 7C,D**), indicating an insufficient supplement of progenitors and retard repopulation in the thymus. In line with decreased ETP and DP cells, the number of single positive CD4^+^ and CD8^+^ T cells also decreased in these mice (**Figure 7E,F**). Furthermore, the naïve T cells in the spleen also showed significant defect in CV1 blockade group (**Figure 7G,H**), while B cell population was normal (**Figure 7I**). Thus, these data suggest that CD47-SIRPα signaling blockade impedes thymic regeneration of T cells.

**Figure 7.**
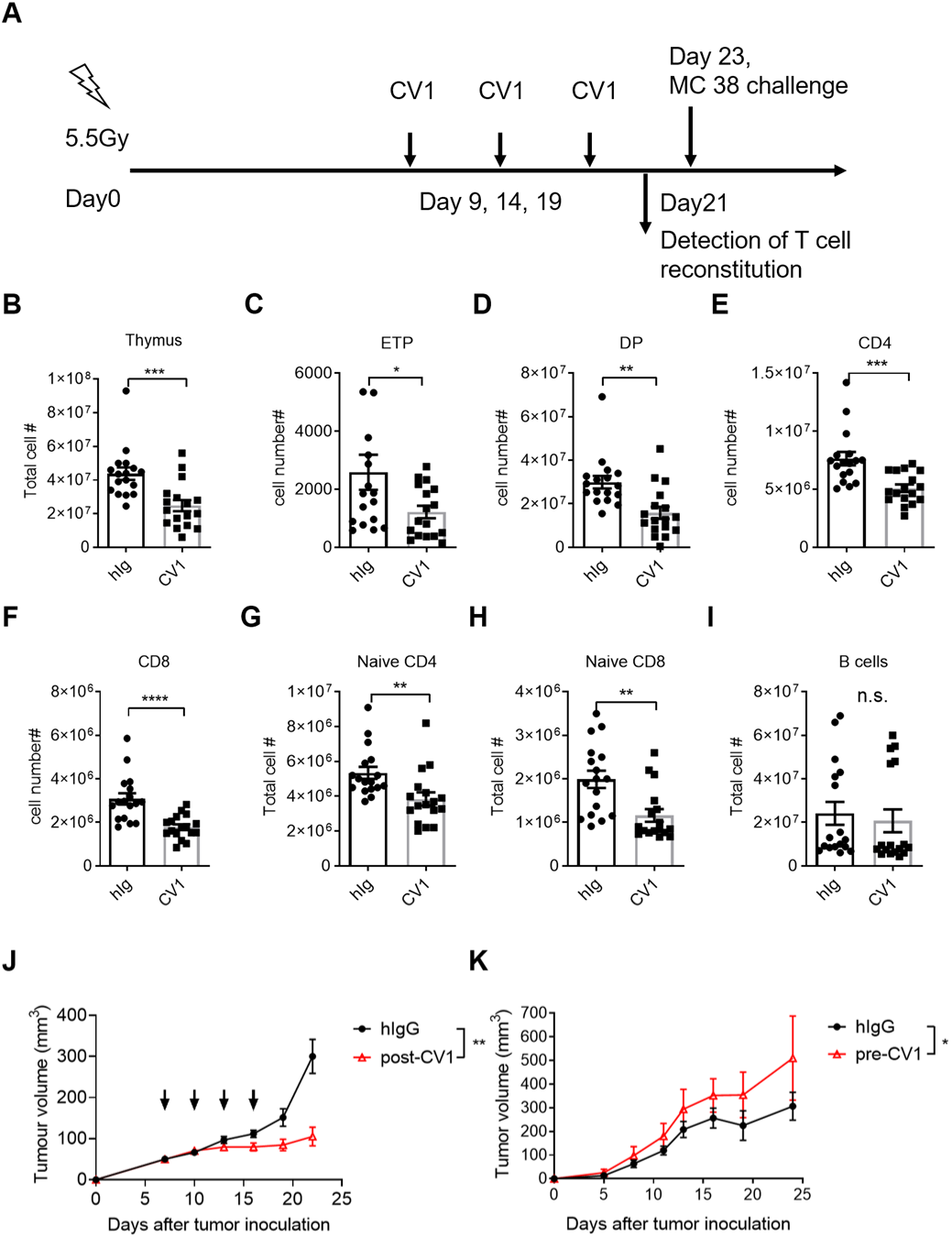
SIRPα signal blockade impairs T cell regeneration and antitumor response upon CRT. (**A**) CV1 or control hIg (200μg/mouse) was injected intraperitoneally into C57BL/6 mice every five days starting from day 9 after sublethal total body irradiation (SL-TBI). In tests related to (**B-I**), mice were sacrificed at day 21 after SL-TBI, in test of (K), mice were challenged with 5*10^5^ MC38 tumors subcutaneously at day 21 after SL-TBI. (**B-F**) The number of total thymocytes (**B**) and indicated subset of ETP (**C**), double positive thymocytes (**D**), CD4^+^ thymocytes (**E**) and CD8^+^ thymocytes (**F**) were analyzed at day 21 as described in (**A**). (**G-I**) The number of CD62L^+^CD44^-^ naïve CD4^+^ T cells (**G**), CD62L^+^CD44^-^ naïve CD8+ T cells (**H**) and B cells (**I**) in the spleen were analyzed at day 21 as described in (**A**). (**J**) Anti-tumor effect of CV1: C57BL/6 mice were subcutaneously injected with 5*10^5^ MC38 tumors, 7 days after tumor inoculation, mice were treated with CV1 or hIgG (50ug/mouse, intra-tumor administration) every 3 days for 4 times (indicated by arrowheads). Tumor growth were monitored and plotted. (**K**) MC38 tumor growth in mice pre-treated with CV1 or control hIgG in the phase of T cell reconstitution, as described in (**A**). Error bars represent s.e.m. Asterisks mark statistically significant difference, \**P*<0.05, \*\**P*<0.01, \*\*\**P*<0.001, \*\*\*\**P*<0.0001, determined by two-tailed unpaired Student’s *t*-test.

As CD47-SIRPα blockade is currently a promising therapeutic antitumor strategy, impaired peripheral T cell regeneration after CV1 treatment led us to concern that SIRPα blockade might dampen T cell-mediated antitumor response during CRT. We therefore challenged the mice experienced SL-TBI plus CV1 treatment with an immunoresponsive murine colon tumor model MC38 (**Figure 7A**). In stark contrast to the fact that CV-1 treatment inhibits established tumor growth in non-SL-TBI mice (**Figure 7J**), a significant faster tumor growth rate was observed in mice experienced SL-TBI and CV1-treatment (**Figure 7K**). These data call attention to the use of CD47-SIRPα blockade reagent for tumor therapy, especially together with CRT.

## Discussion

Homing of bone marrow-derived progenitors to the thymus is the prerequisite of continuous T cell development. In our previous study, Ly6C^-^P-selectin^+^ portal endothelial cells, TPECs(Shi et al., 2016), were found to be the entry gate of HPCs to the thymus. In current study, we have further extended the knowledge about how TPECs interact with progenitors and regulate their thymic entry.

SIRPα has been well recognized as a “don’t eat me” receptor on myeloid cells. Interestingly, our current study reveals a previously totally unrecognized function of SIRPα on ECs for TEM regulation. SIRPα-CD47 interaction has been reported to regulate TEM of neutrophils and monocytes(de Vries et al., 2002; Liu et al., 2002; Stefanidakis et al., 2008). In these studies, CD47, but not SIRPα, is considered as the receptor mediating signal transduction, while SIRPα expressed on neutrophils or monocytes is the ligand. SIRPα engagement of CD47 on epithelial or endothelial cells activates Gi-protein dependent pathway to facilitate transmigration or Rho family GTPase dependent pathway to reorganize EC cytoskeleton. However, in our current work, we clearly showed that SIRPα downstream signal in ECs is required for VE-cadherin endocytosis and TEM. First, in bone marrow chimeric mice that were deficient in hematopoietic SIRPα, thymic ETP is normally maintained (**Figure 2—figure supplement 2E,F**), while non-hematopoietic deficiency of SIRPα resulted in impaired thymic ETPs and thymic progenitor homing (**Figure 2G,H and Figure 2I,J**). These results exclude the possibility in the conventional view that SIRPα is derived from the migrating cells while CD47 is from endothelial compartment. Second, in the absence of functional intracellular region of SIRPα, while keeping extracellular region intact, *Sirpα*-ΔICD or *Sirpα*-Y4F EC monolayer showed deficiency of TEM comparable to that of *Sirpα*^-/-^ ECs (**Figure 5A,B**), underlying the importance of EC-SIRPα downstream signal in controlling TEM. This was further confirmed by the fact that replenishment of WT *Sirpα* but not mutant *Sirpα*-Y4F can rescue TEM reduction in *Sirpα*^-/-^ ECs (**Figure 5B**). Third, *Sirpα*-ΔICD ECs also had impaired endocytosis of VE-cadherin similar to that of *Sirpα*^-/-^ ECs (**Figure 6D**), further supporting the role of EC-SIRPα downstream signaling in TEM.

The role of CD47 in this scenario might be complicated. CD47 is ubiquitously expressed in almost all cells, including immune cells and endothelial cells both of which are involved in this trafficking process. Our data demonstrated that ETP expressed the highest level of CD47 than other subsets of hematopoietic cells during T cell development (**Figure 4A**). The expression level of CD47 on ETPs is also significantly higher than that on TPECs (**Figure 4—figure supplement 1E,F**). These data suggest that migrating cell-derived CD47 might play a major and active role activating SIRPα on ECs for TEM. In fact, CD47 deficient or blocked lymphocytes transmigrate across ECs less efficiently compared to WT control cells when endothelial CD47 is absent (**Figure 4E,F**). Whether physiologically low level of endothelial CD47 constantly work for TEM remains intriguing. Although lymphocyte transmigrate through WT endothelial monolayer more efficiently than CD47 deficient endothelial monolayer (**Figure 4—figure supplement 1H**), given the artificially hyperexpression of CD47 on immortalized WT MS1 cells, this does not necessarily reflect the physiological role of endothelial CD47 on thymic homing. More physiological experimental model is required for further elucidation.

Our study reveals a novel mechanism of adherens junction VE-cadherin endocytosis. Both SHP2 and Src kinase have been reported to regulate VE-cadherin endocytosis, via modification of VE-cadherin in different manners(Allingham et al., 2007; Wessel et al., 2014). However, how they are activated remains unclear. Here, our data suggest that CD47-SIRPα might be one of the upstream signals. In our study, deletion of SIRPα or its cytoplasmic domain results in significant deficiency of TEM and VE-cadherin endocytosis **(Figure 5A and Figure 6A-D**). Regeneration of constitutively activated SHP2 largely rescues TEM in *Sirpα*^-/-^ ECs (**Figure 5C**), while regeneration of ITIM motif mutant SIRPα fails to do so (**Figure 5B**). Together with the fact that ITIM motif activates cytosolic SHP2 phosphatase(Motegi et al., 2003; Tsuda et al., 1998), these data suggest SIRPα signal induced VE-cadherin endocytosis is at least in part through ITIM-SHP2 pathway. In addition, we have also found CD47 engagement induces Src activation in ECs (**Figure 5G,H**). Inhibition of Src kinase via PP2 results in significantly impaired TEM and VE-cadherin endocytosis in WT ECs, but not in *Sirpα*^-/-^ ECs (**Figure 5J and Figure 6E**). Therefore, Src kinase is also involved in SIRPα signal induced VE-cadherin endocytosis. How SIRPα coordinates both SHP2 and Src pathways remains to be investigated in future.

The novel role of SIRPα on TEM regulation might not be limited to TPECs. It is interesting to find that elevated expression level of SIRPα is also found on high endothelial cells (HECs) (data not shown). HECs are specialized blood vascular cells in lymph nodes (LNs). HECs play an important role for LN homing of several immune cell subsets, including naïve T and B cells, plasmacytoid DCs (pDCs) and the precursors of conventional DCs. Examination of peripheral lymph nodes of *Sirpα*^-/-^ mice at resting state revealed no significant change in the percentage/number of these subsets of immune cells (data not shown). Whether SIRPα, expressed on HECs, might regulate immune cell trafficking during certain immune responses remains an intriguing question for future study.

CRT has been commonly used either alone or in combination with immunotherapy for cancer patients(Manukian et al., 2019; Teng et al., 2015). Thymic injury is often observed during CRT(Penit and Ezine, 1989; Zlotoff et al., 2011). Thymic dependent T-lineage regeneration might be important for successful tumor therapy(Joao et al., 2006; Parkman et al., 2006; Politikos and Boussiotis, 2014; Savani et al., 2006). CD47-SIRPα signaling blockade is a promising strategy for tumor immunotherapy and is also combined with CRT(Feng et al., 2019). In our study, we specifically tested the impact of CD47-SIRPα blockade post thymic injury on thymic and peripheral T cell regeneration. Indeed, post SL-TBI induced thymic injury, significantly impaired thymic ETP population and T cell regeneration, both in thymus and at periphery, were found upon CD47-SIRPα blockade. This is also associated with faster growth of MC38 tumors. Whether thymic HPC homing and T cell regeneration is the sole mechanism of CD47-SIRPα blockade for increased tumor growth is still unclear. Nevertheless, our current data provide compelling evidence that post-SL-TBI CD47-SIRPα blockade results in impaired thymic HPC homing and T cell regeneration, and raise a concern about the side effect of CD47-SIRPα blockade on antitumor immunity, especially in combination with CRT.

In summary, we have discovered a novel function of thymic endothelial SIRPα in controlling thymic progenitor homing, revealed its underlying molecular mechanism in regulating adherens junctional VE-cadherin endocytosis. In addition, our data suggest the importance of SIRPα-mediated thymic and peripheral T cell regeneration in the context of tumor therapy.

## Materials and Methods

### Mice

WT C57BL/6 mice were purchased from Vital River, a Charles River company in China; Tie2-Cre mice were purchased from Nanjing Biomedical Research Institute (Nanjing, China). *Sirpα^flox/flox^* mice were kindly provided by Dr. Hisashi Umemori (Children’s Hospital, Harvard Medical School, Boston, MA); *Sirpα^-/-^* mice were kindly provided by Dr. Hongliang Li (Wuhan University, China); *Cd47*^-/-^ mice were gift from Dr. Yong-guang Yang (The First Bethune Hospital of Jilin University, China). All mice are on C57BL/6 background and were maintained under specific pathogen-free (SPF) condition. Mice for experiments are four to eight weeks old and sex-matched unless otherwise specified. Animal experiments were performed according to approved protocols of the institutional committee of the Institute of Biophysics, Chinese Academy of Sciences.

### Cell lines

Mouse pancreatic islet endothelial cell line MS1 were passaged in DMEM (Hyclone) supplemented with 10% FBS (BI) and 1% penicillin-streptomycin (Gibco).

1. *Sirpα^-/-^* and *Cd47^-/-^* MS1 were constructed by transfecting pLentiCRISPR v2 (addgene, Feng Zhang, #52961) cloned with oligos for desired single guide RNA. Two pairs of oligos for Sirpα targeting were designed for in case one of them fails to work.

Sirpα-1-F: caccGCAGCGGCCCTAGGCGGCCA,
Sirpα-1-R: aaacTGGCCGCCTAGGGCCGCTGC;
Sirpα-2-F: caccGCCCGGCCCCTGGCCGCCTA
Sirpα-2-R: aaacTAGGCGGCCAGGGGCCGGGC;
2. MS1 cell line with truncated *Sirpα* lacking intracellular domain (*Sirpα*-ΔICD) was constructed by transfecting pLentiCRISPR v2 cloned with oligos targeting exon 6 of the first cytoplasmic region from N-terminus. The resulting clones were sequenced for +/− 1 frameshift, which created frame shift and advanced stop codon within exon 7. Sirpα ITIM motifs are within exon 8.

Sirpα-ΔICD-E6-1-F: caccGAGGGGTCAACATCTTCCACA,
Sirpα-ΔICD-E6-1-R: aaacTGTGGAAGATGTTGACCCCTC;
3. Sirpα-4F and Sirpα-WT overexpressing lines were constructed based on *Sirpα*^-/-^ MS1. Full-length Sirpα coding sequence was cloned from WT MS1 cells, and point mutated at all four tyrosine (Y) position from TAC or TAT to TTC or TTT respectively, thus become non-functional phenylalanine (F). Cloning primers for Sirpα CDS:

XbaI_Sirpα-F: ATATTCTAGACC ACCATGGAGCCCGCCGGCCCG,
Sirpα_BamHI-R: TATGGATCCTCACTTCCTCTGGACCTGGA. Wild type or mutated (4F) form of Sirpα were ligated to pTK643-GFP with multiple cloning site (MCS) and packaged for lentivirus. MS1 cells were infected for 24 hours with polybrene before seeded for monoclonal selection. FACS analysis of GFP and SIRPα surface staining was used to identify overexpressed clones.
4. Shp2-E76K overexpressing line was constructed based on *Sirpα*^-/-^ MS1. Full-length Shp2 coding sequence was cloned from mouse genome and point mutated at Glutamic Acid 76 (E76) to constitutively active Lysine (K). Shp2-E76K was then ligated to pTK643-GFP and *Sirpα*^-/-^ MS1 was infected as above described. Cloning primer for Shp2 CDS:

XbaI-Shp2-F: ATTtctagaGCCACCatgACATCGCGGAGATGG,
XbaI-Shp2-R: cgtctagaaTCATCTGAAACTCCTC TGCT

### Isolation of thymic ECs

Thymus was collected, digested and Percoll enriched as previously described(Shi et al., 2016). Briefly, thymus was digested in RPMI 1640 medium with 2% fetal bovine serum, 0.2 mg/ml collagenase I, 1U/ml dispase and 62.5μg/ml DNase for four rounds of 20-minute digestion on a 37℃, 120rpm shaker. Dissociated cells after each round and the final digest were washed and applied for discontinuous Percoll gradient enrichment.

Cells were resuspended with 1.115 g/ml Percoll at the bottom, and 1.065 g/ml Percoll and PBS were laid on the middle and top layer, respectively. After being centrifuged at 2700rpm (872g) at 4℃ for 30 minutes, EC-containing cells from the upper interface were collected and subjected to subsequent analysis.

### Cell preparation

Bone marrow cells were isolated from mouse femurs and tibias. In brief, soft tissues were cleared off and the ends of the bones were cut, bone marrow was than flushed out using a 23-gauge needle containing ice-cold PBS. Whole-tissue suspensions of thymus, lymph node and spleen were generated by gently forcing the tissue through a 70μm cell strainer. Red blood cells were lysed with ACK in samples of bone marrow and spleen.

### Flow cytometry and cell sorting

Flow cytometry data were acquired with a LSRFortessa cell analyzer (BD Biosciences) and analyzed using FlowJo software (BD Biosciences). Cell sorting was performed on a FACSAria III cell sorter (BD Biosciences). The following fluorescent dye-conjugated antibodies against cellular antigens were used for: 1) analysis of the thymic portal endothelia cells (TPECs): CD45 (30-F11), CD31 (MEC13.3), P-selectin (RB40.34) and Ly-6C (HK1.4); SIRPα (P84) 2) analysis of the lineage negative progenitor cells: CD45.1 (A20), CD45.2 (104), Lineage cocktail: CD11b (M1/70), CD11c (N418), NK1.1 (PK136), Gr-1 (RB6-8C5), B220 (RA3-6B2) and TER-119 (Ter-119); 3) analysis of the early t cell progenitors (ETPs): Lineage cocktail (see above), CD4 (GK1.5), CD8 (53-6.7), CD25 (PC61), CD44 (IM7) and c-Kit (2B8); 4) analysis of Lin^-^Sca-1^+^c-Kit^+^ (LSK) and Common lymphoid progenitors (CLPs): Lineage cocktail (see above), c-Kit (2B8), Sca-1 (D7), IL7Rα (A7R34) and Flt3 (A2F10); 5) analysis of peripheral lymphoid subsets: CD4 (GK1.5), CD8 (53-6.7), B220 (RA3-6B2), CD62L (MEL-14), CD44 (IM7). 6) Other antibodies used: CD47 (miap301), VE-cadherin (BV13), VCAM-1 (429). 7) Corresponding isotypes: Rat IgG1 κ iso (RTK2071), Rat IgG2a κ iso (eBR2a). Dead cells were excluded through DAPI (Sigma-Aldrich) or LIVE/DEAD (ThermoFisher) positive staining.

### Quantitative real-time PCR

Thymic endothelial cells were gated as subset I (P-selectin^-^Ly-6C^+^), subset II (P-selectin^+^Ly-6C^+^) and subset III, which is TPEC (P-selectin^+^Ly-6C^-^). These three subsets were FACS sorted and RNA was extracted using RNeasy Micro Kit (Qiagen). The quality and quantity of total RNA was assessed using a Nanodrop 2000c spectrometer (Thermo Scientific). cDNA was synthesized using RevertAid First Stand cNDA Synthesis Kit (Thermo) and Oligo (dT)_18_ primer. Gene expression was quantified using the following primers. Quantitative real-time PCR was performed using SYMBR Premix Ex Taq (Takara) and run on Applied Biosystems 7500 Real-Time PCR System. Relative mRNA expression was calculated with a standard curve and C_T_ value of target gene and *β-actin* or *Gapdh* control.

Sirpα-F: TGCTACCCACAACTGGAATG,
Sirpα-R: CCCTTGGCTTTCTTCTGTTT,
mActb-F: ACACCCGCCACCAGTTCGC,
mActb-R: ATGGGGTACTTCAGGGTCAGG;
mGapdh-F: AACCACGAGAAATATGACAACTCACT,
mGapdh-R: GGCATGGACTGTGGTCATGA.

### SIRPα-hIg (CV1) production and SIRPα signal blockade

SIRPα-hIg was produced as previously described(Liu et al., 2015). Briefly, pEE12.4-CV1, kindly provided by Dr. Yang-Xin Fu (University of Texas Southwestern Medical Center, Dallas, USA), was transiently expressed in FreeStyle 293 expression system (Thermo Fisher). SIRPα-hIg was then purified with Sepharose Protein A/G beads. In progenitor short-term homing assay, a single dose of 200μg (2μg/ml) CV1 or control hIgG (Sigma) was administrated intraperitoneally (i.p.) 2 days before congenic bone marrow adoptive transfer. In thymic regeneration assay, 200μg CV1 or control hIgG was administrated i.p. every five days since day 9 after SL-TBI.

### Bone marrow chimeras and short-term homing assay

6 weeks old recipient mice received 10 Gy γ radiation once. 5×10^6^ donor bone marrow cells were injected intravenously into the recipients on the day or the next day of irradiation. 8 weeks later, mice are ready for subsequent studies. Short-term homing assay: donor bone marrows were taken from femurs and tibias from WT mice, red blood cells were lysed and labeled by 2μM CFSE, and 5×10^7^ bone marrow cells were transferred intravenously to the recipients. The recipients were sacrificed 2 days later and detected for donor derived progenitors in the thymus and spleen.

### Thymic regeneration and blood sampling

Wild type mice were exposed to 5.5 Gy γ radiation once (sub-lethal total body irradiation, SL-TBI) with no exogenous hematopoietic cell infusion. Mice were then maintained in SPF facility for up to 35 days. Recovery of thymic regenerated T-lymphocytes were monitored by blood sampling from orbital sinus every five days since day 9 after SL-TBI. The volume of collected blood was recorded and lymphocyte cell number was counted by FACS.

### Lymphocyte adhesion assay

The adhesion assay was performed mainly as previously described(Au - Lowe and Au - Raj, 2015). 1×10^5^/mL MS1 cells were plated on coverslip (NEST, φ=15mm), growing for 24 hours and then stimulated with 10ng/mL TNFα for additional 24 hours. 1×10^6^ total lymphocytes from axillary and inguinal lymph nodes were added to the top of coverslip and incubated for 3 hours. Dunk in and out the coverslip vertically of the PBS five times before trypsinized and collect total cells for lymphocyte counting by FACS.

### Transmigration assay

2.5×10^4^ MS1 cells were initially plated on Transwell filter (Corning, φ=6.5mm, with 5.0μm pore), allowed growing for 24 hours and then stimulated with 10ng/mL TNFα for additional 24 hours before transmigration assay. 1×10^6^ total lymphocytes were added per Transwell. Chemoattraction was achieved by adding 10ng/mL CCL19 (PeproTech) in the bottom chamber. Four hours later, migrated lymphocytes were collected for cell counting and subset analysis by FACS.

### VE-cadherin endocytosis assay

The endocytosis assay was performed mainly as previously described(Wessel et al., 2014). In this assay, 1×10^5^/mL MS1 cells were plated on coverslip, allowed growing for 24 hours and then preactivated with 10ng/mL TNFα for additional 24 hours. Cells grown to confluency were treated for 1 hour with 150μM chloroquine prior to the endocytosis assay. Cell were then incubated for 30 minutes at 37 ℃ with anti-VE-cadherin-biotin (BV13) in culture medium. Antibody was then washed and 1×10^6^ total lymphocytes were added to the top of coverslip. After 1 hour incubation at 37 ℃, lymphocytes were removed by quickly rinsing wells with prewarmed culture medium. And surface-bound antibodies were removed by washed for 3 times, 20 seconds each time, with acidic PBS (pH 2.7 PBS with 25mM glycine and 2% FBS). Cells were then fixed with 1% paraformaldehyde in PBS for 5 minutes followed by permeabilization with 0.5% Triton-x-100 for 10 minutes at room temperature. Internalized primary antibodies were then detected by fluorescence conjugated Streptavidin (Biolegend). DNA was stained with DAPI. Five or more fields were observed in each sample on a Zeiss LSM 700 confocal system (63×/1.4 Oil) with locked parameters. Endocytosed VE-cadherin was quantified via home-made script running under ImageJ batch mode with fixed signal threshold for each experiment. Threshold value was adjusted according to signal background from isotype control sample.

### Western blotting

MS1 cells were lysed with lysis buffer containing 20mM Tris·HCl, 2mM EDTA, 0.5% NP-40, 1mM NaF and 1mM Na_3_VO_4_ and protease inhibitor cocktail. The samples were heated to 95℃ for 5 minutes with loading buffer containing 0.5% 2-Mercaptoethanol.

Equal amount of samples was loaded and resolved on a 10% SDS-PAGE gel. Proteins were then transferred to PVDF membrane (Millipore, 0.45μm). The membranes were first incubated with anti-phospho-Src (Tyr416) antibody (CST) or anti-phospho-Src (Tyr529) antibody (Abcam) followed by Goat anti-Rabbit HRP (CWBIO, China). The blot was developed by chemiluminescent HRP substrate (ECL, Millipore). The blot was then stripped and reblotted with anti-β-actin antibody (Zsbio, China) subsequently. The images were captured on a Tanon 5200 chemi-image system (Tanon, China). Gel images were quantified with Lane 1D analysis software (SageCreation, China).

### VE-cadherin real-time imaging

Flow chamber was designed and made by Center for Biological Imaging (CBI), Institute of Biophysics (IBP), Chinese Academy of Sciences (CAS). Briefly, a coverslip (NEST, φ=25mm) with endothelial monolayer can be inserted into the thermostatic (37℃) flow chamber and supplied with pre-warmed flow medium (DMEM). Flow medium was controlled by a pump (adjusted by frequency and voltage). The flow chamber was then fixed to an adaptor which allowed the bottom side of the coverslip fit into observation range of the 60× immersion oil lens of an Olympus FV1200 spectral inverted laser scanning confocal microscopy. MS1 endothelial cells were plated on the coverslip sited in 6-well plate at a density of 1.2×10^5^/mL and activated by 10ng/mL of TNFα 24 hours later. After additional 24 hours, MS1 endothelial cells were sequentially labeled with rat-anti-mouse VE-cadherin, anti-Rat-Alexa Fluor 488 and anti-CD31-Alexa Fluor 647. CD4^+^ T cells were prepared from mouse inguinal and axillary lymph nodes, and divided into two parts: one part blocked by CD47 antagonist CV1 at 10ug/mL and labeled by anti-CD4-V450, another part controlled by hIg and labeled by anti-CD4-PE. Two parts of CD4^+^ T cells were equally mixed and resuspend to 1×10^6^/mL in flow medium. Focus was limited to the layer that maximized junctional CD31 signal, and Z-axis drift compensation (ZDC) was activated to lock focus. Continuous multi-channel imaging on AF488, PE, AF647 and V450 was conducted and concatenated. Colocalization of VE-cadherin with CD31 at sites of differentially treated T cells was measured and calculated by Imaris 9 software. In reciprocally labeling tests (**Figure 6—figure supplement 1d,e**), CV1-treated T cells were labeled with anti-CD4-PE, and hIg-treated T cells were labeled with anti-CD4-V450.

### RNA-Seq and microarray data analysis

The RNA-Seq data of thymic ECs was previously published and available as GSE_83114(Shi et al., 2016). Differentially expressed genes of TPECs (fold change>2 and p value of pairwise t-test <0.01 in either the contrast of TPECs versus subset II or subset I EC subset) were filtered for their function by gene ontology term cell migration (GO_0016477). The hints were candidates for further analysis. Microarray data from previously published work(Lee et al., 2014) were analyzed by Affymetrix Expression Console (1.4.1) and Transcriptome Analysis Console (3.0) software following manufacturer’s instruction. Z-score normalized heatmap was generated by gplots package in R (3.6).

### Statistical analysis

Statistical analyses were performed using GraphPad 6.0 (Prism). Two-tailed unpaired Student’s t-test was used for significance test, unless otherwise specified. All experiments with significant differences were performed independently three times. Experiments with no significant differences were performed at least two times. The results were expressed as the mean ± s.e.m. (n.s., not significant; *P < 0.05; **P < 0.01; ***P < 0.001; and****P < 0.0001).

## Acknowledgements

We would like to thank Dr. Hisashi Umemori (Children’s Hospital, Harvard Medical School, USA) for providing *Sirpα^flox/flox^* mice, Dr. Hongliang Li (Wuhan University, China) for *Sirpα^-/-^* mice, Dr. Yong-guang Yang (The First Bethune Hospital of Jilin University, China) for *Cd47*^-/-^ mice and Dr. Yang-Xin Fu (University of Texas Southwestern Medical Center, USA) for pEE12.4-CV1 expression plasmid. We would like to thank Xiaoyan Wang, and Yihui Xu in the Key Laboratory of Infection and Immunity, Institute of Biophysics (IBP), Chinese Academy of Sciences (CAS) for providing instrumental support on flow cytometry and confocal imaging; Yan Teng, Yun Fen in the Center for Biological Imaging, IBP, CAS for their innovative instrument for real-time confocal imaging in flow chamber. This work was supported by grants from National Natural Science Foundation of China (31770959 and 82025015 to M.Z.).

## Author contributions

B.R. and M.Z. designed the experiments and analyzed the data; B.R. conducted most experiments with some help from H.X., Y.L., Z.W., and Y.S. B.R. and M.Z. wrote the manuscript; M.Z. supervised the study.

## Competing interests

The authors declare no competing interests.

---

**Figure 2—figure supplement 1.**
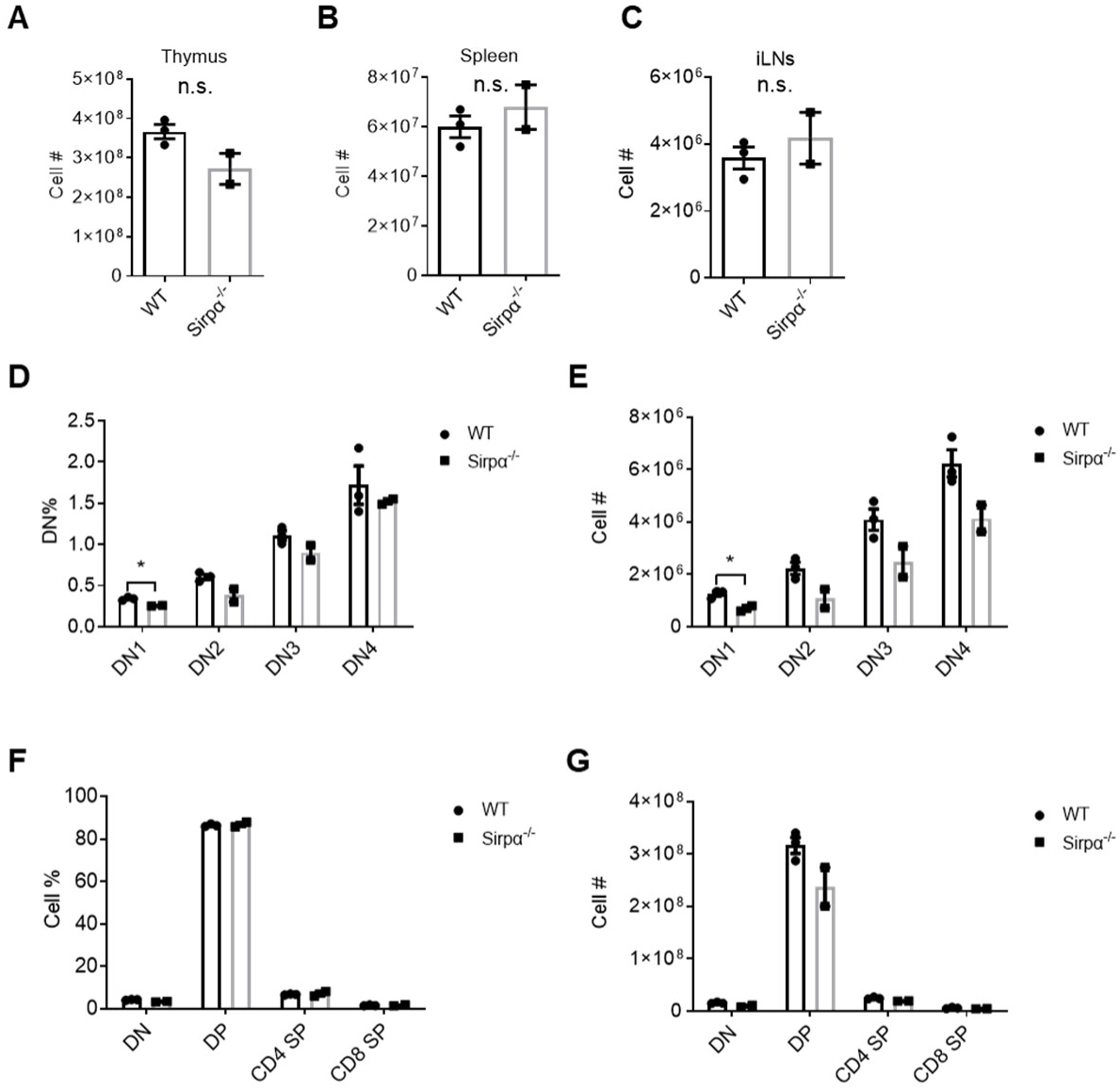
SIRPα deficiency does not alter major T cell development in resting state. (**A-E**) Cellularity of the thymus (**A**), spleen (**B**) and bilateral inguinal lymph nodes (**C**), proportion (**D**) and total cell number (**E**) of double negative thymocyte subsets (DN1, DN2, DN3 and DN4). (**F,G**) proportion (**F**) and total cell number (**G**) of major thymocytes subsets (DN, double negative, DP, double positive, CD4 SP, CD4 single positive and CD8 SP, CD8 single positive) among total thymocytes in *Sirpα^-/-^* and control mice. Error bars represent s.e.m. Asterisks mark statistically significant difference, \**P*<0.05, n.s. not significant, determined by two-tailed unpaired Student’s *t*-test.

**Figure 2—figure supplement 2.**
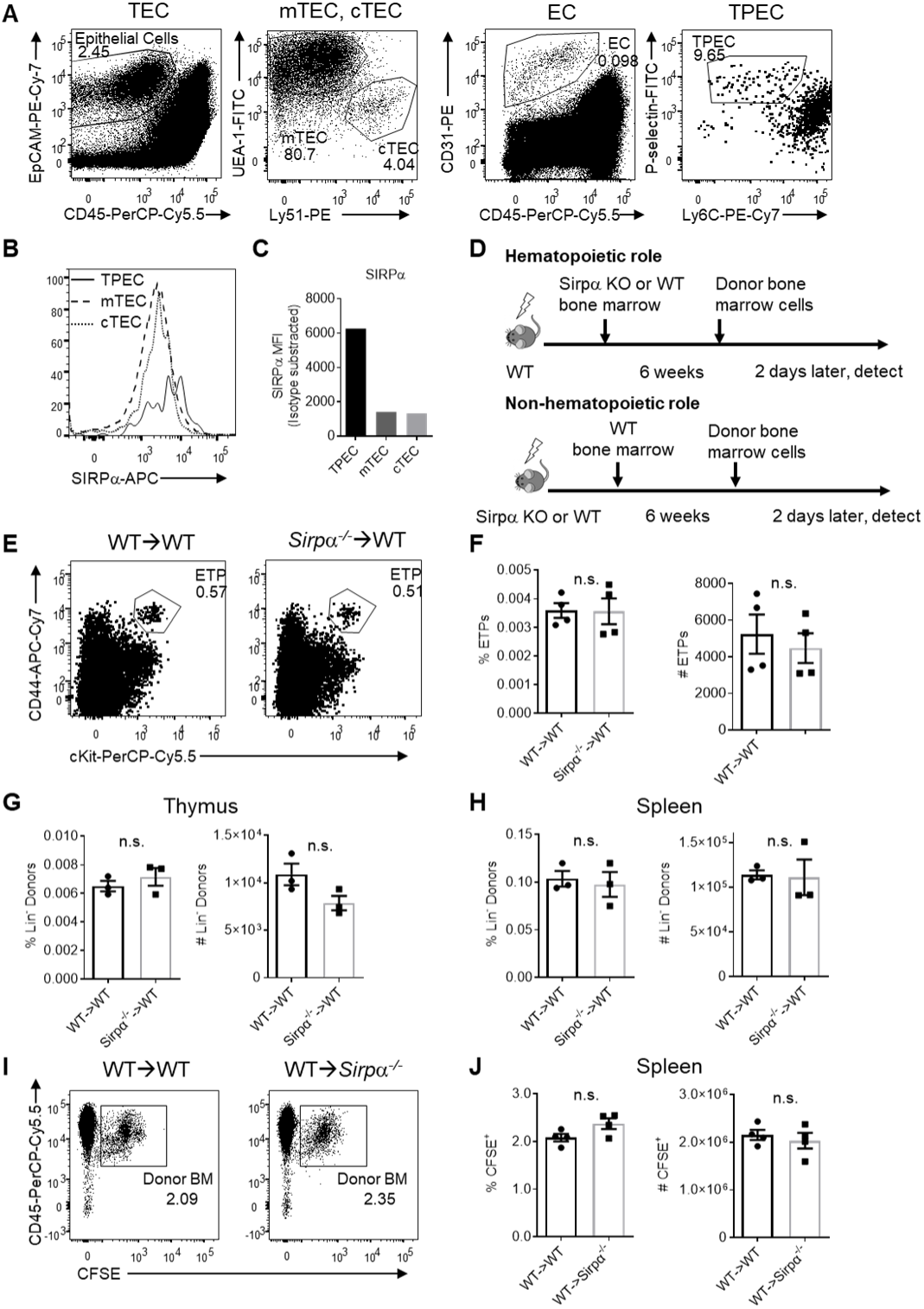
Endothelial SIRPα is essential for thymic progenitor homing. (**A**) Representative dot plot of thymic epithelial cell (TEC, EpCAM^+^CD45^-^), TEC subsets (mTEC, UEA-1^+^Ly51^-^ and cTEC, UEA-1^-^Ly51^+^), thymic endothelial cells (EC, CD31^+^CD45^-^) and EC subsets (TPEC, P-selectin^+^Ly6C^-^). (**B,C**) Expression (**B**) and statistics (**C**) of SIRPα on TPEC, mTEC and cTEC subsets of thymic stroma. (**D**) Schematic diagram showing thymic progenitor short-term homing assay in bone marrow chimeric mice. (**E,F**) Flow cytometric analysis (**E**) and statistics (**F**) of proportion and absolute cell number of ETP population in WT mice reconstituted with *Sirpα^-/-^* or WT bone marrows. (**G,H**) Short-term homing assay in WT mice reconstituted with WT or *Sirpα^-/-^* bone marrows. Lineage^-^ donor progenitor cells were determined in the thymus (**G**) and in the spleen (**H**). (**I,J**) Short-term homing assay in WT or Sirpα^-/-^ mice reconstituted with WT bone marrows. Total donor cells (CFSE^+^CD45^+^) in the spleen were determined. Error bars represent s.e.m. n.s. means not significant, determined by two-tailed unpaired Student’s *t*-test.

**Figure 3—figure supplement 1.**
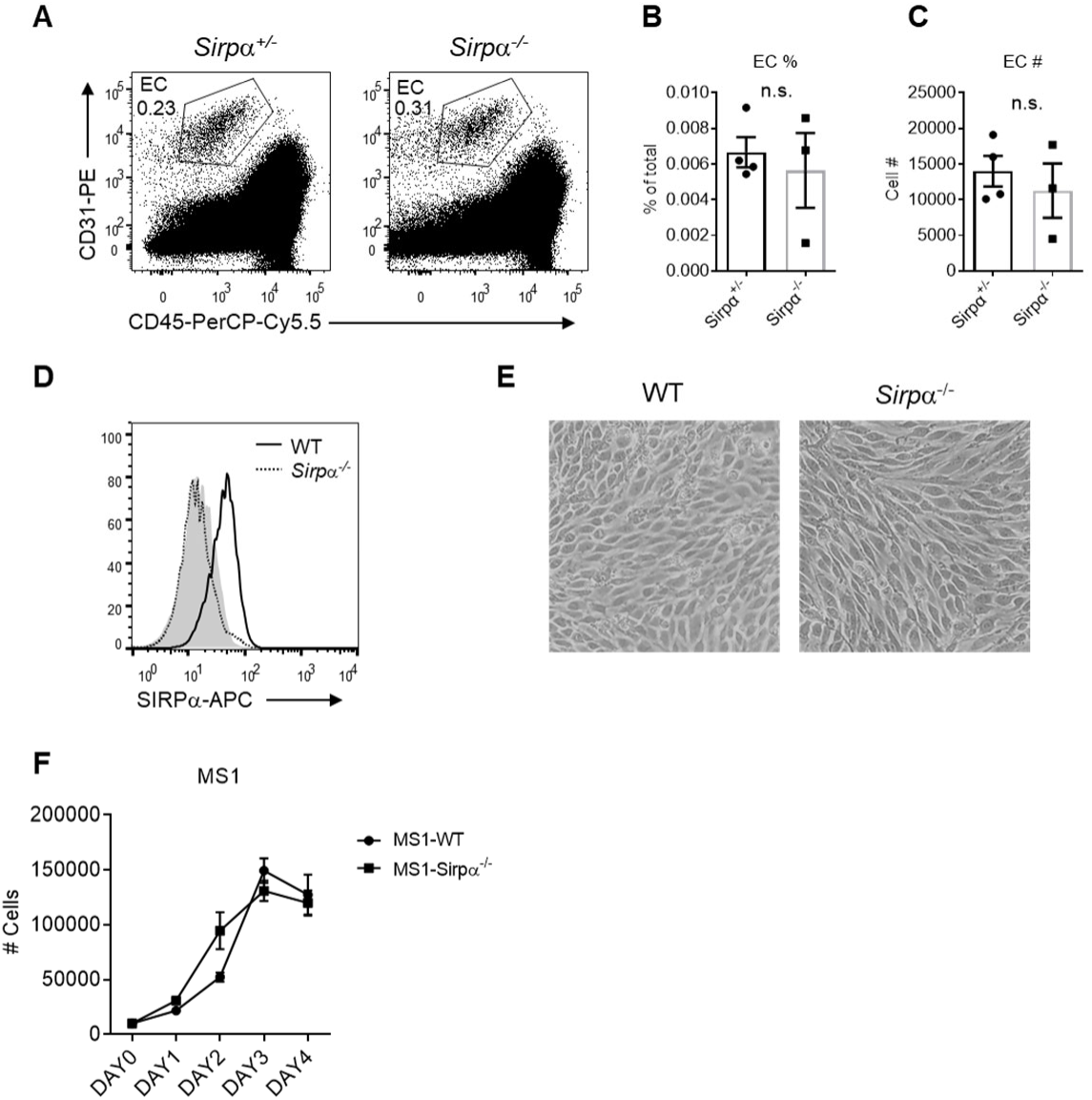
SIRPα does not control EC development and growth. (**A**) Representative flow cytometric analysis of thymic endothelial cells (EC, CD31^+^CD45^-^) in the thymus of *Sirpα^-/-^* and control mice. (**B,C**) Proportion (**B**) and corresponding cell number (**C**) of ECs in total thymocytes in a thymus. (**D**) SIRPα detection by flow cytometry in CRISPR/Cas-9 mediated *Sirpα^-/-^* MS1 cells, shadow indicates isotype level. (**E**) Microscopic observation (4×) of MS1 cells. (**F**) FACS analysis of MS1 cell number in the transwell chamber at indicated time points. Error bars represent s.e.m. n.s. means not significant, determined by two-tailed unpaired Student’s *t*-test.

**Figure 4—figure supplement 1.**
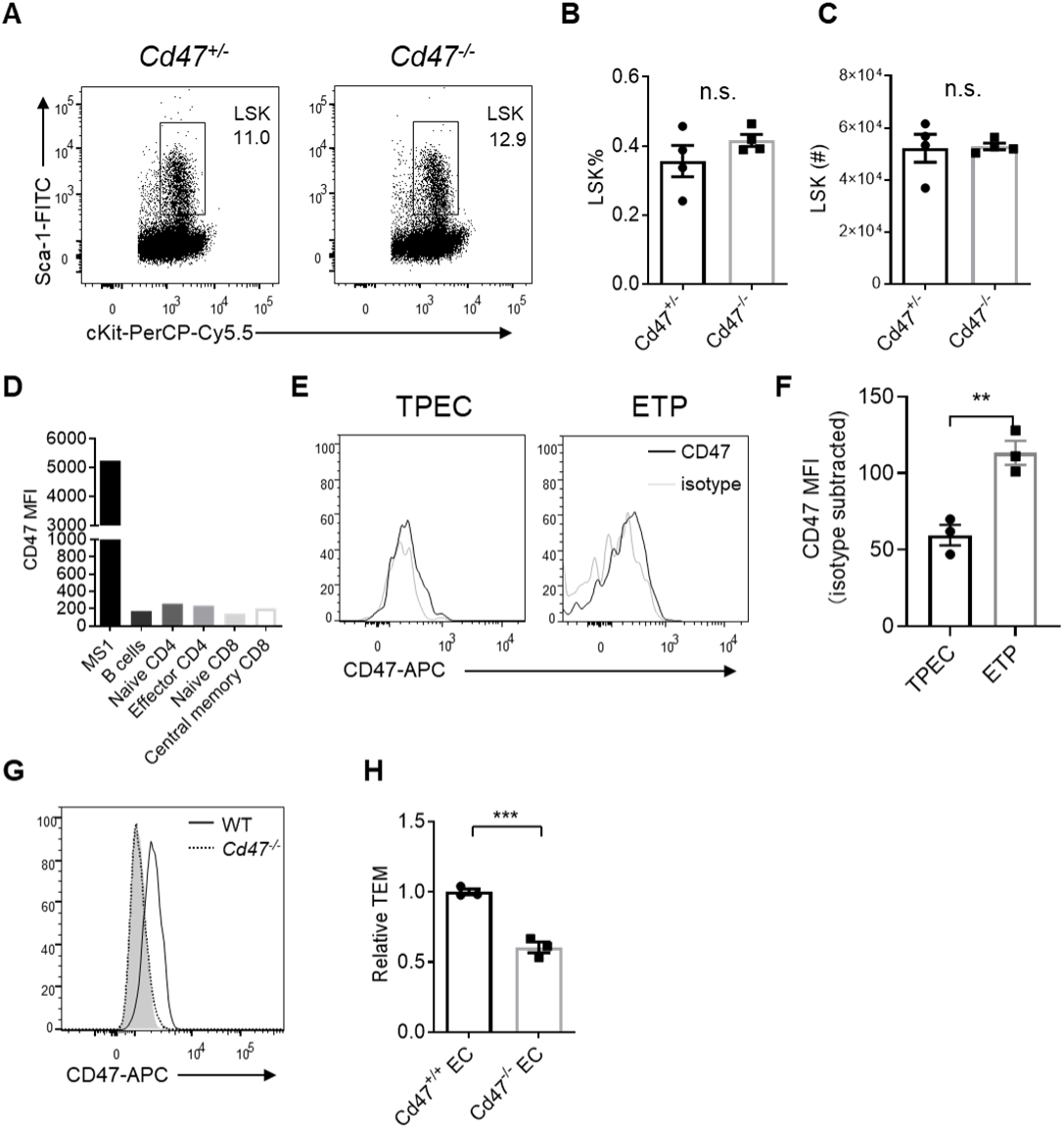
Migrating cell-derived CD47 guides their TEM. (**A**) Flow cytometric analysis of LSK progenitor cells (Lineage^-^cKit^+^Sca-1^+^) in the bone marrow of *Cd47^-/-^* and control mice. (**B,C**) Proportion (**B**) and corresponding cell number (**C**) of LSKs of total bone marrow cells in a pair of femurs and tibias. (**D**) CD47 expression on MS1 endothelial cells and primary lymphocyte subsets from inguinal lymph nodes. B cells (B220^+^), naïve CD4 (CD4^+^CD62L^+^CD44^-^), effector CD4 (CD4^+^CD62L^-^CD44^+^), naïve CD8 (CD8^+^CD62L^+^CD44^-^) and central memory CD8 (CD8^+^CD62L^+^CD44^+^) T cells. (**E**) CD47 expression on TPEC and ETP subsets (Lineage^-^CD25^-^CD44^+^cKit^+^). (**F**) Statistics of CD47 expression on TPECs and ETPs. (**G**) CD47 detection by flow cytometry in CRISPR/Cas-9 mediated *Cd47^-/-^* MS1 cells, shadow indicates isotype level. (**H**) Lymphocyte transmigration through *Cd47*^-/-^ or WT (*Cd47*^+/+^) MS1 endothelial monolayer. Error bars represent s.e.m. Asterisks mark statistically significant difference, \**P*<0.05, \*\**P*<0.01, \*\*\**P*<0.001, n.s. not significant, determined by two-tailed unpaired Student’s *t*-test. Raw data of transmigration assay in H are available in SupplementaryFile1.

**Figure 5—figure supplement 1.**
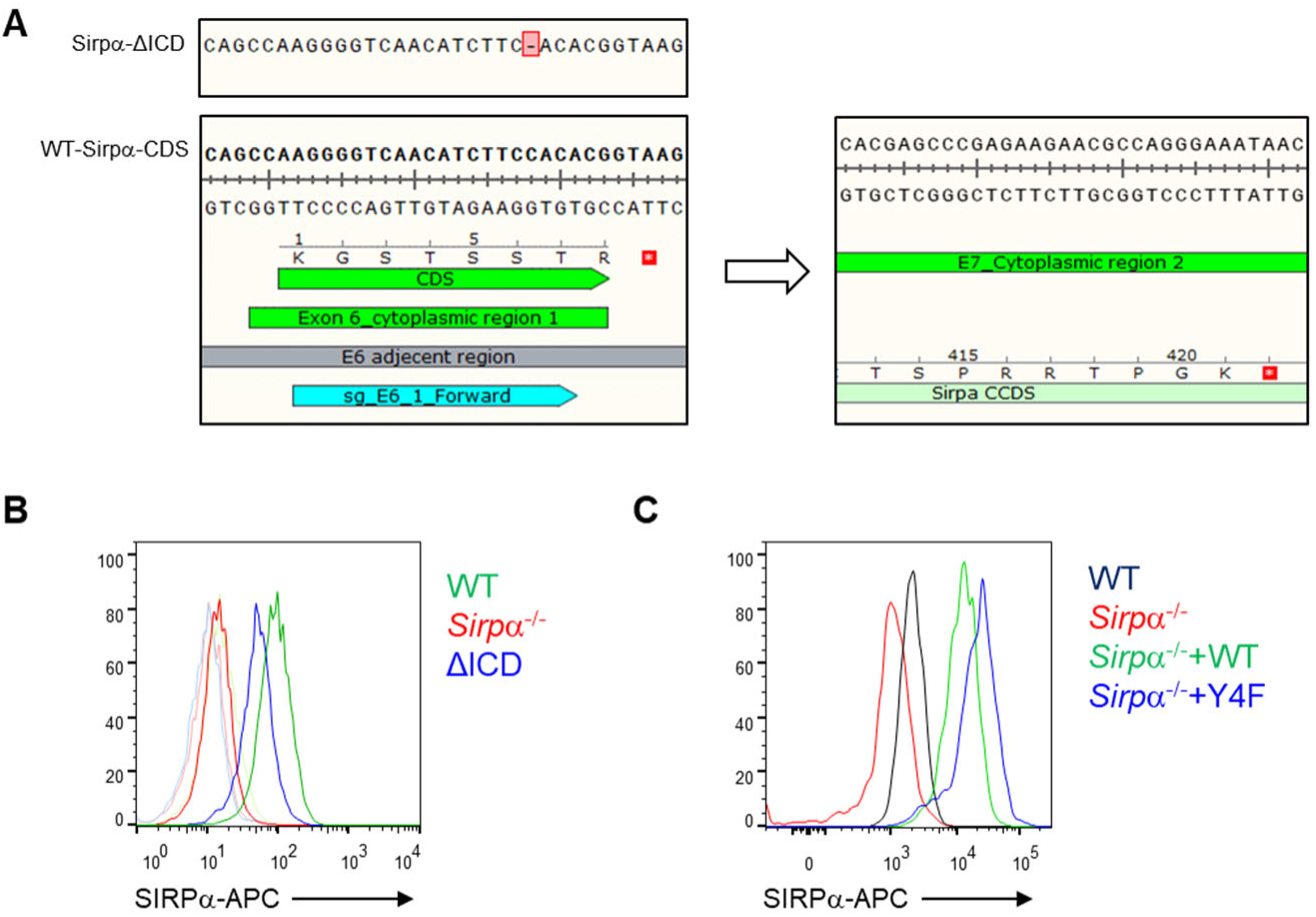
Construction of SIRPα mutant MS1 cell lines. (**A**) Schematic view of Cas9 mediated truncation of *Sirpα* intracellular region. sgRNA targeting Exon 6 of *Sirpα*, the first exon encoding for the cytoplasmic region, was designed to create frame-shift mutation at the exon which generated stop codon TAA (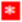) at the following exon 7, which terminates SIRPα translation before intracellular signal region within exon 8 to form a presumably truncated form of intracellular region of SIRPα (SIRPα-ΔICD). (**B**) FACS analysis of surface expression of SIRPα on WT, intracellular domain truncated (ΔICD) and *Sirpα*^-/-^ MS1 cell lines, tint lines indicate corresponding isotype level. (**C**) FACS analysis of SIRPα expression after lentiviral-mediated overexpression of WT coding sequence of *Sirpα* (*+Sirpα* WT) or *Sirpα* with intracellular tyrosine residues mutated to phenylalanine (+*Sirpα*-4F) which inactivated intracellular signal of SIRPα, on the basis of *Sirpα^-/-^* MS1.

**Figure 6—figure supplement 1.**
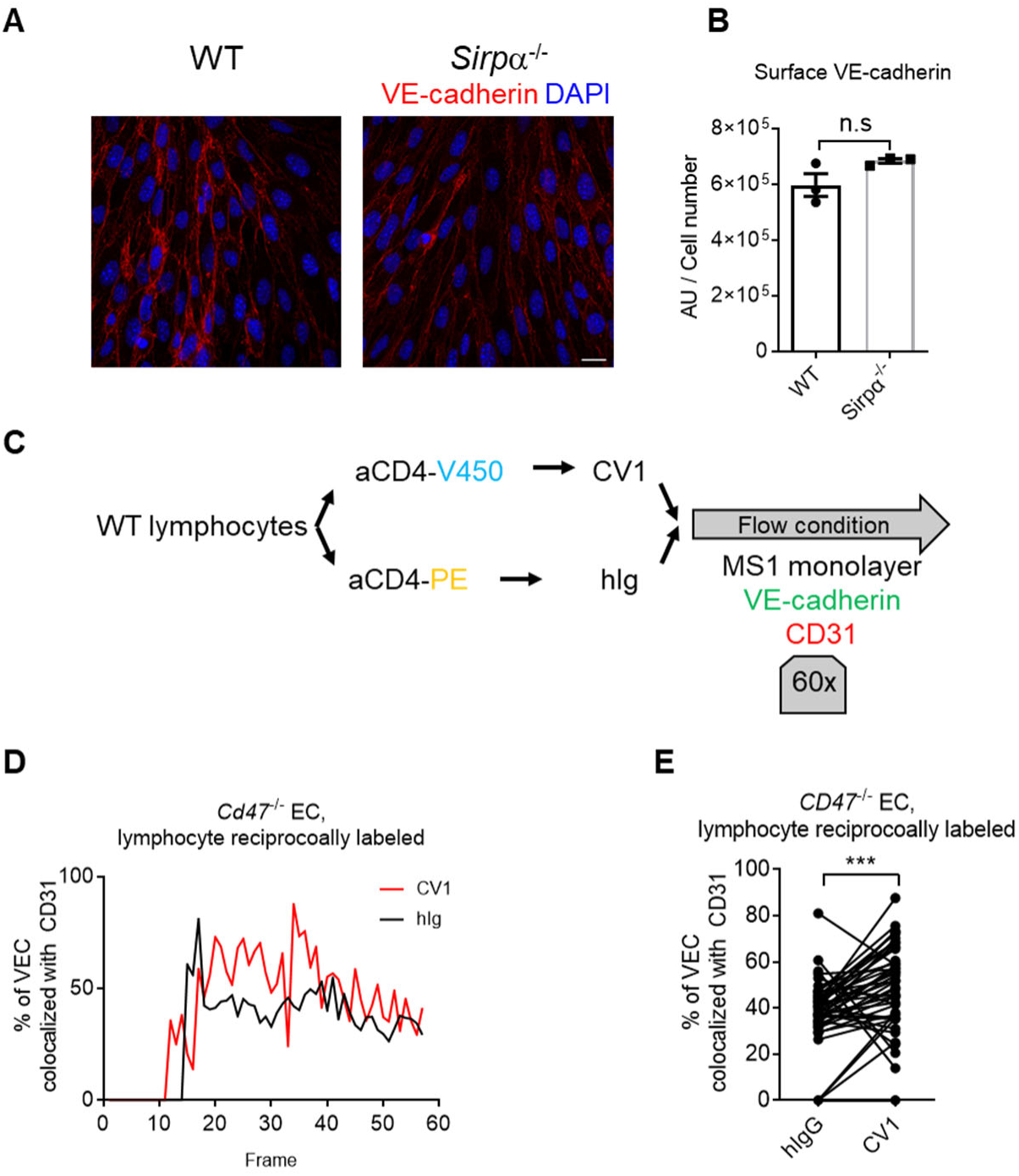
CD47-SIRPα signaling promotes VE-cadherin endocytosis. (**A**) Confocal microscopy of distribution of VE-cadherin on WT and *Sirpα* KO MS1 monolayers, bar represents 20μm. (**B**) VE-cadherin signal per cell calculated by total VE-cadherin signal intensity divided by cell number counted by DAPI highlighted nucleus. Significance determined by two-tailed unpaired Student’s *t*-test. (**C**) Schematic view of real-time analysis of adherens junctional VE-cadherin in the presence of migrating lymphocytes. (**D,E**) Measurement (**D**) and statistical analysis (**E**) of VE-cadherin colocalization with CD31 after CD4^+^ T cell injection, CV1- and control hIg-treated CD4^+^ T cells are reciprocally labeled by fluorescent dyes as used in Figure 6F,G. Significance determined by two-tailed paired Student’s t-test. Error bars represent s.e.m. Asterisks mark statistically significant difference, \*\*\**P*<0.001, n.s. not significant. Statistics of real time imaging in D and E are available in SupplementaryFile1.

## Notes

### Competing Interest Statement

The authors have declared no competing interest.

